# High-Throughput Screening of the Effects of 90 Xenobiotics on the Simplified Human Gut Microbiota Model (SIHUMIx): A Metaproteomic and Metabolomic Study

**DOI:** 10.1101/2023.12.04.569900

**Authors:** Victor Castañeda-Monsalve, Laura-Fabienne Fröhlich, Sven-Bastiaan Haange, Masun Nabhan Homsi, Ulrike Rolle-Kampczyk, Qiuguo Fu, Martin von Bergen, Nico Jehmlich

## Abstract

The human gut microbiota is a complex microbial community with critical functions for the host, including the transformation of various chemicals. While effects on microorganisms has been evaluated using single-species models, their functional effects within more complex microbial communities remain unclear. In this study, we investigated the response of a simplified human gut microbiota model (SIHUMIx) cultivated in an in vitro bioreactor system in combination with 96 deep-well plates after exposure to 90 different xenobiotics, comprising 54 plant protection products and 36 food additives and dyes, at environmentally relevant concentrations. We employed metaproteomics and metabolomics to evaluate changes in bacterial abundances, the production of Short Chain Fatty Acids (SCFAs), and the regulation of metabolic pathways.

Our findings unveiled significant changes induced by 23 out of 54 plant protection products and 28 out of 36 food additives across all three categories assessed. Notable highlights include azoxystrobin, fluroxypyr, and ethoxyquin causing a substantial reduction (log_2_FC <-0.5) in the concentrations of the primary SCFAs: acetate, butyrate, and propionate. Several food additives had significant effects on the relative abundances of bacterial species; for example, acid orange 7 and saccharin led to a 75% decrease in *Clostridium butyricum*, with saccharin causing an additional 2.5-fold increase in *E. coli* compared to the control. Furthermore, both groups exhibited up- and down-regulation of various pathways, including those related to the metabolism of amino acids such as histidine, valine, leucine, and isoleucine, as well as bacterial secretion systems and energy pathways like starch, sucrose, butanoate, and pyruvate metabolism.

This research introduces an efficient in vitro technique that enables high-throughput screening of the structure and function of a simplified and well-defined human gut microbiota model against 90 chemicals using metaproteomics and metabolomics. We believe this approach will be instrumental in characterizing chemical-microbiota interactions especially important for regulatory chemical risk assessments.

## 1 Introduction

The human intestine harbors trillions of microorganisms that play crucial roles in host health and disease. This community, known as the human microbiome, has emerged as a pivotal determinant of host health and well-being (The Integrative HMP (iHMP) Research Network Consortium, 2014; Hajiagha et al., 2021); however, so far, it has not been considered in the hazard assessment of xenobiotics. This is due to the lack of reproducible and reliable model systems and difficulties in establishing causal relationships. Thus, it is necessary to develop reliable screening tools for xenobiotics that detect how the microbiome and its metabolites that are involved in microbiome-host interactions are affected.

The extensive capacity of metabolizing enzymes within the microbiome not only enables the conversion of complex carbohydrates and the production of short-chain fatty acids (SCFAs) that contribute to overall nutrition and immune regulation (Rodríguez-Carrio et al., 2017) but also extends to the transformation of xenobiotics. Xenobiotics are substances that are taken voluntarily, such as pharmaceuticals (Zimmermann-Kogadeeva et al., 2020), as well as semi-voluntarily taken supplements, such as dietary supplements (Cao et al., 2020), and non-intentionally consumed chemicals (Popli et al., 2022), such as plant protection products in contaminated food and water (Elmassry et al., 2022). Given that approximately 80% of chemical exposure occurs through the digestive tract, its effects on the microbiome and microbial biotransformation are becoming increasingly important.

Following their interactions with the microbiota, ingested xenobiotics can lead to a wide range of outcomes. These include metabolite activation, inactivation, transformation into a toxic form, or the absence of biotransformation while exerting an antimicrobial effect on specific species or the entire microbiota (Spanogiannopoulos et al., 2016). These interactions have the potential to give rise to the generation of altered microbial metabolites, accumulation of specific substances, and modifications in the structural composition of the community (Lichtman et al., 2016). These outcomes may subsequently culminate in potentially modified microbiota with corresponding changes in its functional attributes (Chen and Sundrud, 2018).

The disruption of host-associated microbes due to exposure to external chemicals is a well-known phenomenon. However, it is important to note that our understanding of xenobiotic metabolism in gut microbes is limited. Metabolic functions rarely correlate directly with microbial phylogeny and there is considerable strain-level variation, even within the same species. Large-scale metagenomic analyses have not provided experimental evidence to support predicted changes in xenobiotic metabolic activities (Koppel et al., 2017). Until this inquiry is addressed, regulatory agencies are unlikely to incorporate microbiota into their risk assessment strategies, even though they recognize their potential future significance from both toxicokinetic and toxicodynamic perspectives (Ampatzoglou et al., 2022). Consequently, additional research is needed to elucidate the specific gut microbes, genes, and enzymes involved in xenobiotic metabolism, and their biological effects.

Numerous strategies have emerged to improve the analysis of microbial communities and effectively complement and expand the traditional genomic profiling methodologies. These approaches encompass the exploration of data types that accurately portray the functional dynamics of microbiota, such as metatranscriptomics, metaproteomics, and metabolomics. This paradigm shift has yielded refined mechanistic models that elucidate both the structural composition and operational aspects of microbial communities (Franzosa et al., 2015). Metaproteomics has emerged as a valuable tool for precisely delineating functional dimensions, including translation, energy metabolism, carbohydrate utilization, and antimicrobial defense mechanisms (Wissenbach et al., 2016). Identifying proteins not only allows taxonomic assignment but also facilitates the attribution of functional properties. Therefore, metaproteomics is an optimal methodology for exhaustively investigating the multifaceted functional repertoire of gut microbiota (Haange and Jehmlich, 2016; Lohmann et al., 2020).

In this context, there are metaproteomic-based screening tools described that employ batch cultures of stool donors to assess the effects of xenobiotics on the composition and function of intestinal microbial communities (Li et al., 2019a). Short-term batch cultures are relatively easy to manage and yield valuable insights into the metabolism of xenobiotics as well as their impact on the structure and function of microbial communities (Li et al., 2019b). However, considerable biological variance among stool donors limits reproducibility (Zhong et al., 2019). Moreover, the complexity of these consortia restricts the depth of mechanistic insights possible, and a brief incubation time resembles acute intoxication more than a chronic one.

Continuous cultivation of a more representative model system of the intestinal microbiome, such as OligoMM (Brugiroux et al., 2016) or SIHUMIx (Becker et al., 2011), can be employed to address these shortcomings. The latter encompasses approximately 90% of the biochemical functions of the microbiome and allows cultivation periods of up to 21 d (Schäpe et al., 2019). Another advantage is its high reproducibility and the availability of well-established procedures for proteomic and metabolomic analyses (Schäpe et al., 2020; Petruschke et al., 2021).

In this study, we make a significant contribution to the ongoing efforts aimed at deepening our understanding of the mechanisms underlying microbiome-mediated toxicity. To achieve this goal, we systematically examined the impact of 90 distinct xenobiotics, including 54 plant protection products and 36 food additives and dyes, on a simplified model of the human microbiome (hereafter referred to as SIHUMIx). Importantly, our approach involved the utilization of xenobiotic concentrations that aligned with the legally permissible levels established for these compounds. We gained a comprehensive understanding of SIHUMIx’s response to these chemical compounds by observing modifications in its structural dynamics and functionality via metaproteomic analyses. Furthermore, we quantitatively evaluated the abundance of SCFAs using metabolomics.

## 2 Materials and Methods

### 2.1 Simplified Human Intestinal Microbiota Model – SIHUMIx

In this experiment, the extended simplified human intestinal microbiota (SIHUMIx) (Becker et al., 2011) was utilized, which comprised a total of eight species: *Anaerostipes caccae* (DSMZ 14662), *Bacteroides thetaiotaomicron* (DSMZ 2079), *Bifidobacterium longum* (NCC 2705), *Blautia producta* (DSMZ 2950), *Clostridium butyricum* (DSMZ 10702), *Erysipelatoclostridium ramosum* (DSMZ 1402), *Escherichia coli* K-12 (MG1655), and *Lactobacillus plantarum* (DSMZ 20174).

### 2.2 *In vitro* bioreactor cultivation and xenobiotic concentrations

Prior to inoculation, each strain was thawed in 9 mL of Brain Heart Infusion (BHI) broth under anaerobic conditions with shaking at 175 rpm and a temperature of 37°C. Overnight cultures were counted, and 1×10^9^ cells per strain were used, resulting in an inoculum of 8×10^9^ cells per bioreactor. Three 250 mL Multifors 2 bioreactors (Infors, Switzerland) filled with sterile Complex Intestinal Media (CIM) (**Supplementary Table S1**) were inoculated with SIHUMIx in batch mode. After 24 h, the bioreactors were transitioned to continuous cultivation at a media exchange rate of 24 h. Previous studies have shown that during the initial four days after inoculation, SIHUMIx undergoes a stabilization phase until day 5, when the community reaches a stationary state characterized by stable species relative abundance and metabolic outputs (Krause et al., 2020).

### 2.3 Determination of xenobiotic concentration and exposure of SIHUMIx to 90 chemicals in deep-well plates

A total of 54 plant protection products (**Supplementary Table S2**) and 36 food additives and dyes (**Supplementary Table S3**) were used in this study. The concentration of each chemical was set at 0.5% of its maximum recommended intake value, such as Acceptable Daily Intake (ADI), No Observed Adverse Effect Level (NOAEL), or Average Dietary Exposure (More et al., 2021). To determine this concentration, we considered that the average human stool contained approximately 1 × 10^11^ bacterial cells per gram (Stephen and Cummings, 1980), and the average daily human stool production was 200 g. Based on these data, it was estimated that approximately 2 × 10^13^ bacterial cells can be exposed to 100% of the recommended intake of each chemical. To reflect a relevant exposure scenario, the concentration was determined to be 0.5% of the maximum recommended intake value owing to the proportion of bacterial cells used in this study compared to the average fecal content.

Aliquots of SIHUMIx were harvested from each bioreactor and pooled under anaerobic conditions every day, from day 6 to day 12, for a total of seven days. We used a modified version of the RapidAIM model (Li et al., 2019a). Within an anaerobic chamber (Coy Laboratory Products, Grass Lake, MI, USA), 100 µL of SIHUMIx were inoculated into 890 µL of Complex Intestinal Media (CIM), along with 10 µL of the chemical, to maintain a solvent concentration below 2%. Prior to inoculation, both the sterile CIM and chemical stocks were pre-reduced in an anaerobic chamber. This process resulted in a total working volume of 1 mL. This procedure was performed in triplicates for each chemical using 96 deep well plates. To prevent cross-contamination and gas accumulation, the plates were sealed with a silicon mat with a small hole on the top. Incubation was performed at 37°C with continuous shaking (Avantor VWR; Radnor, PA, USA) at 320 rpm for 24 h. In total, we used 14 deep-well plates, with two plates used per day. Additionally, each plate included a negative control consisting of SIHUMIx inoculated in CIM along with the solvent of each chemical. After 24 h, each culture was centrifuged at 5,000x g for 10 min at 4°C. The resulting pellet and supernatant were subsequently frozen at -80°C until further processing (**Figure. 1**).

**Figure 1:**
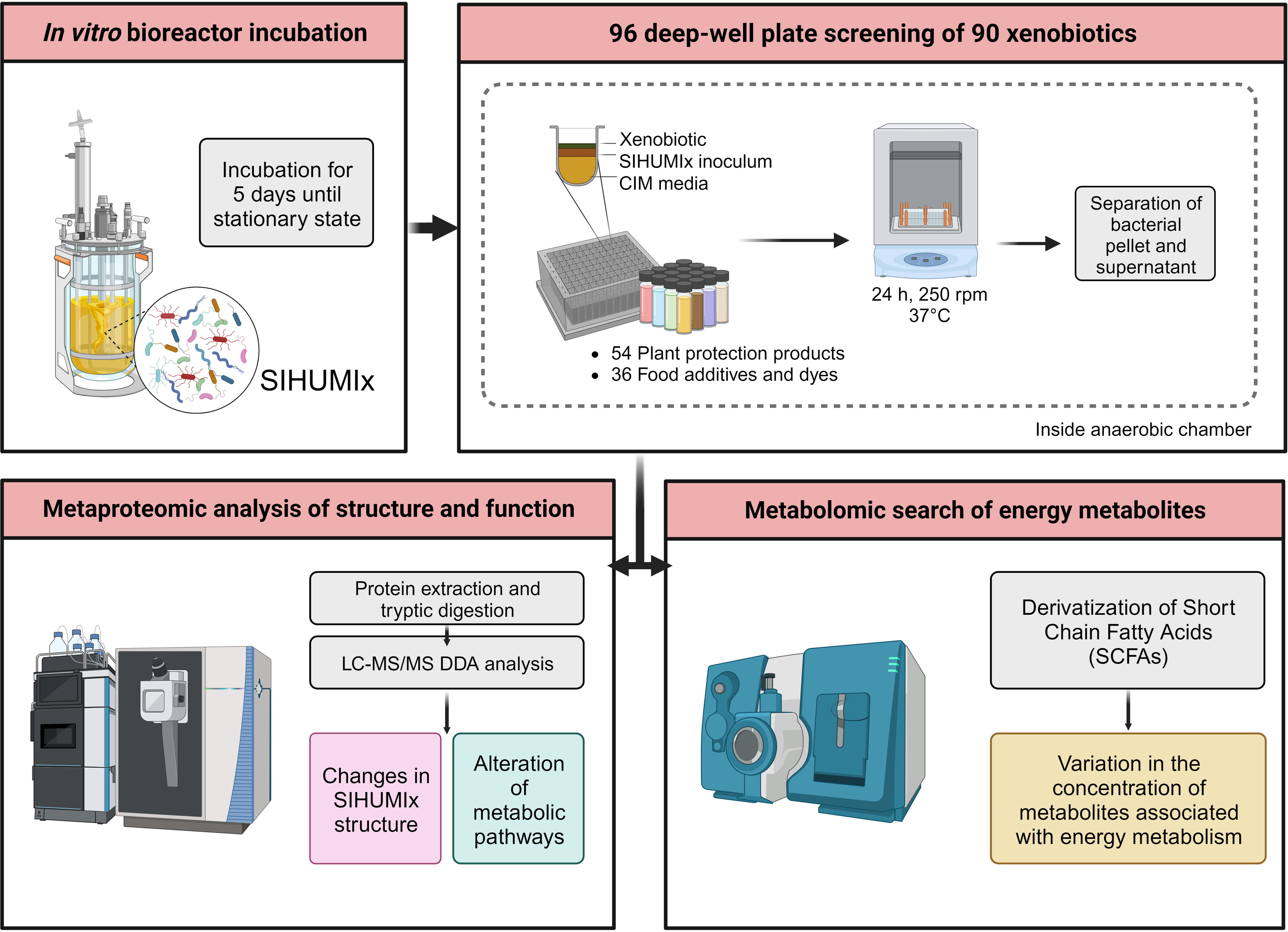
Experimental overview of the screening of 90 xenobiotics with the simplified model of human microbiota (SIHUMIx). The community was inoculated in bioreactors until steady-state. Then, SIHUMIx were cultured in 96-deep-well plates at 37°C under anaerobic conditions for 24h exposed to the individual xenobiotics. Pellets and supernatants were collected for metaproteomic and metabolomic analysis respectively.

To ensure quality throughout the seven-day screening process, each day underwent testing with both positive and negative controls for bacterial metabolism. The positive control involved exposure of SIHUMIx to Sulfasalazine (CAS 599-79-1), a drug commonly used for irritable bowel disease. This drug relies on bacterial azoreductases to cleave the molecule and release its active components (Yaqoob et al., 2022). The positive control mimicked chemical exposure inoculation, replacing the xenobiotic with sulfasalazine at a final concentration of 1µM. Conversely, the negative control contained sulfasalazine and CIM, without the addition of SIHUMIx.

After every day of the screening, bacterial metabolism control samples were thawed, vortexed, and centrifuged (10 000 rpm, 4°C, 10 minutes). In order to precipitate the macromolecules in the media, supernatants were diluted 1:1 in cooled methanol (-20°C) and centrifuged as in the previous step. For analysis, the cleared supernatants were diluted with water 1:100 and 10 µL injected for analysis. Quantification of Sulfasalazine was performed via RPLC-MS/MS on an Agilent 1260 Infinity series HPLC system (Agilent Technologies, Waldbronn, Germany), an Atlantis T3 column, 2.1 × 100 mm (Waters Corp., Milford, USA) with a SecurityGuard C18 guard column (4 × 2.0 mm2; Phenomenex), connected to a QTRAP 6500 (Sciex, Darmstadt, Germany) mass spectrometer. Chromatographic separation was performed at a flow rate of 0.4 mL/min and 35°C, using mobile phases A1 water and B1 methanol, each containing 2mM ammonium formate and 0.1% acetic acid. The gradient of A1 decreased from 90% after 1 minute to 45% after 6 minutes, then to 5% after 19 minutes, and then to 5% until 23 minutes. Lastly, gradient was returned to starting conditions and flushed until 29 minutes. Measurement was performed in MRM and ESI negative with a mass transition from *m*/*z* 397 to 197 (quantifier) and 397 to 240 (qualifier). MultiQuant 3.0.3 (Sciex, Framingham, US) was used to quantify Sulfasalazine via external calibration.

Only the days that showed a reduction below limit of detection of the Sulfasalazine in the biotic controls were used for metaproteomic and metabolomic analysis.

### 2.4 Metaproteome analysis

Samples were prepared as previously described (Riesbeck et al., 2022) with some modifications. Briefly, preserved pellets of SIHUMIx were resuspended in a lysis buffer consisting of 8M urea, 2M thiourea, and 1 mM phenylmethylsulfonyl fluoride. To disrupt the bacterial cells, bead beating was performed using the FastPrep-24 system (MP Biomedicals, Santa Ana, CA, USA; 5.5 ms, 1 min, 3 cycles), followed by heat shock at 90°C for 10 min with 1,400 rpm (ThermoMixerTM, Eppendorf, Germany) and centrifugation (10,000× g, 10 min). The resulting supernatant was used for protein concentration determination using a Pierce 660 nm Protein Assay (Thermo Fisher Scientific, Waltham, MA, USA).

Vivacon 500 columns with a 10 kDA Molecular weight cutoff membrane (Sartorius, Göttingen, Germany) were equilibrated with lysis buffer and centrifuged (14,000× g, 20°C, 20 min), and the remainder of the centrifugation steps were performed under the same conditions. Subsequently, 50 µg of the protein was loaded onto the column and centrifuged. The proteins that adhered to the filter were then incubated with 200 µL of 10 mM dithiothreitol in lysis buffer using a ThermoMixer (Eppendorf) at 37°C and 600 rpm for 1 min and then with no agitation for 30 min. After centrifugation, the protein pellets underwent further alkylation and proteolytic cleavage, as described previously (Wiśniewski et al., 2009). Finally, the resulting peptide lysates were dissolved in 0.1% formic acid (FA) for subsequent mass spectrometric measurements.

For each nanoLC-MS measurement, 1 μg of the peptide lysate was injected into a Vanquish Neo nanoHPLC system (Thermo Fisher Scientific). The peptide lysates were initially trapped on a C18-reverse phase trapping column (Acclaim PepMapTM 100, 75 μm × 2 cm, particle size 3 μM, nanoViper, Thermo Fisher Scientific), and then separated on a C18-reverse phase analytical column (Double nanoViper™ PepMap™ Neo, 75 μm × 150 mm, particle size 2 μM, Thermo Fisher Scientific). The separation was achieved using a two-step gradient with mobile phases A (0.01% FA in H2O) and B (80% acetonitrile in H2O and 0.01% FA). During the first step of the gradient, which lasted 95 min, the proportion of mobile phase B was increased from 4% to 30%. This was followed by a 40-minute period in which the proportion of mobile phase B increased from 30% to 55%. The flow rate during the separation was maintained at 300 nL/min.

The eluted peptides were ionized using a Nanospray Flex™ Ion Source (Thermo Fisher Scientific) and detected using an Orbitrap Exploris™ 480 mass spectrometer (Thermo Fisher Scientific). The mass spectrometer was operated with the following settings: for MS, the scan range was set to 350–1,550 *m/z*, the resolution was set to 120,000, the automatic gain control (AGC) target was set to 3’000,000 charges, and the maximum injection time was set to 100 ms. An intensity threshold of 8,000 ions and dynamic exclusion time of 20 s were also applied. The top 10 most intense ions were selected for MS/MS analysis, with an isolation window of 1.4 *m/z*, resolution of 15 000, AGC target of 200 000 ions, and a maximum injection time of 100 ms.

### 2.5 Metaproteome data analysis

Mass spectrometric data processing was conducted using Proteome Discoverer software (v2.5, Thermo Fischer Scientific), employing the SequestHT search engine. The search settings were configured as follows: trypsin (full) enzyme specificity, allowing for a maximum of two missed cleavage sites; precursor mass tolerance of 10 ppm; and fragment mass tolerance of 0.02 Da. The carbamidomethylation of cysteine residues was designated as a fixed modification. To enhance peptide identification, the INFERYS rescoring method was employed, and false discovery rates (FDR) were determined at 1% using Percolator (Gessulat et al., 2019). SequestHT searches were performed using a constructed proteome database, which included the reference proteomes from the eight bacterial strains downloaded from UniProt (www.uniprot.org).

All data quality control, and statistical analyses and correlations were performed using in-house written scripts in R studio (v2023.03.0). Protein functions and pathway assignments were performed using Ghostkoala in KEGG. Only pathways containing a minimum of five proteins and a minimum total coverage of 10% were selected for further analysis.

To assess significant changes in species abundance between the treatment group and the respective control group, protein intensity data was normalized and log-transformed, and an unpaired t-test was performed in GraphPad Prism (v9.4.1 Windows, GraphPad Software, San Diego, California, USA). Additionally, we applied p-value correction using the Holm–Šídák method to account for multiple comparisons.

An ensemble feature selection method, combining the decisions of six feature selection methods (FSMs) that included mean absolute difference, ANOVA F-value, mutual information, minimum redundancy maximum relevance, backward elimination, and random forest, was applied to identify the most relevant subsets of significant differential pathways among the control and treatment groups. A pathway was considered significant if it had an adjusted p-value of less than 0.05 and was ranked in the top 20 pathways by at least four out of the six FSMs. This stringent criterion ensured that only the most robust and consistently identified pathways were significantly affected by the treatment.

### 2.6 Analysis of Short Chain Fatty Acids (SCFA)

The measurement of Short Chain Fatty Acids (SCFA) was conducted following a previously established protocol (Schäpe et al., 2020). The samples were combined with acetonitrile to achieve a final concentration of 50%. To derivatize SCFAs, 0.5 volumes of 200 mM 3-nitrophenylhydrazine and 0.5 volumes of 120 mM N-(3-dimethylaminopropyl)-N-ethylcarbodiimide hydrochloride in pyridine were added to the samples. Derivatization was performed at 40 °C and 300 rpm for 30 min. Subsequently, the derivatized SCFA solutions were diluted 1:50 in a solution containing 10% acetonitrile. Finally, SCFAs were quantified using the diluted solution.

Diluted SCFA derivatives (10 µL) were injected into an RSLC UltiMate 3000^®^ system (Thermo Fisher Scientific) coupled with a QTRAP 5500^®^ mass spectrometer (AB Sciex, Framingham, MA, USA). Chromatographic separation of SCFAs was achieved using an Acquity UPLC BEH C18 column (1.7 µm; Waters, Eschborn, Germany) with water (0.01% formic acid, FA) and acetonitrile (0.01% FA) as mobile phases. The flow rate was set at 0.35 mL/min, and the column temperature was maintained at 40 °C. The elution gradient consisted of 2 min at 15% B, followed by a 15-minute gradient from 15% to 50% B, and a 1-minute hold at 100% B. The column was then equilibrated for 3 min at 15% B. SCFAs were identified and quantified using a scheduled MRM method, with specific transitions for each SCFA. Peak areas were determined using the Analyst® Software (v1.6.2, AB Sciex), and the areas for individual SCFAs were exported.

Quantification of the metabolites was performed using custom R scripts developed in R Studio (v2023.03.0) with calibration curves. Statistical comparisons between each treatment and its respective controls were conducted using unpaired t-tests in GraphPad Prism. To address multiple comparisons, p-value correction was applied using the Holm–Šídák method.

## 3 Results

We investigated the influence of 54 plant protection products and 36 food additives and dyes on the composition of SIHUMIx, which serves as a model for human gut microbiota. All seven days passed the bacterial metabolism control, showing a reduction of sulfasalazine below the limit of detection (**Supplementary Figure S1**). On average, 3,985 distinct protein groups were detected (**Supplementary Table S4**) and a global summary of the results of all treatments that caused a significant change on SIHUMIx’s structure, pathways and SCFA concentrations can be found in (**Supplementary Tables S5 and S6**).

### 3.1 Changes in SIHUMIx community after chemical exposure

First, we analyzed the impact of xenobiotics on community structure. Therefore, the abundance of each bacterial strain of SIHUMIx was assessed by quantifying the protein biomass. Our findings revealed that 14 of the 54 tested plant protection products significantly affected at least one SIHUMIx species (**Figure. 2A**), These products led to changes in the abundance of a particular species, with the log_2_FC ranging from -0.3 to 0.7 when compared to the control. Interestingly, among these 14 plant protection products, only three treatments affected more than one species; fluroxypyr and napropamide affected two, and foramsulfuron affected five. The resulting clusters of plant protection products displayed in **Figure 2A** do not show any trends according to the class or use of xenobiotics.

**Figure 2:**
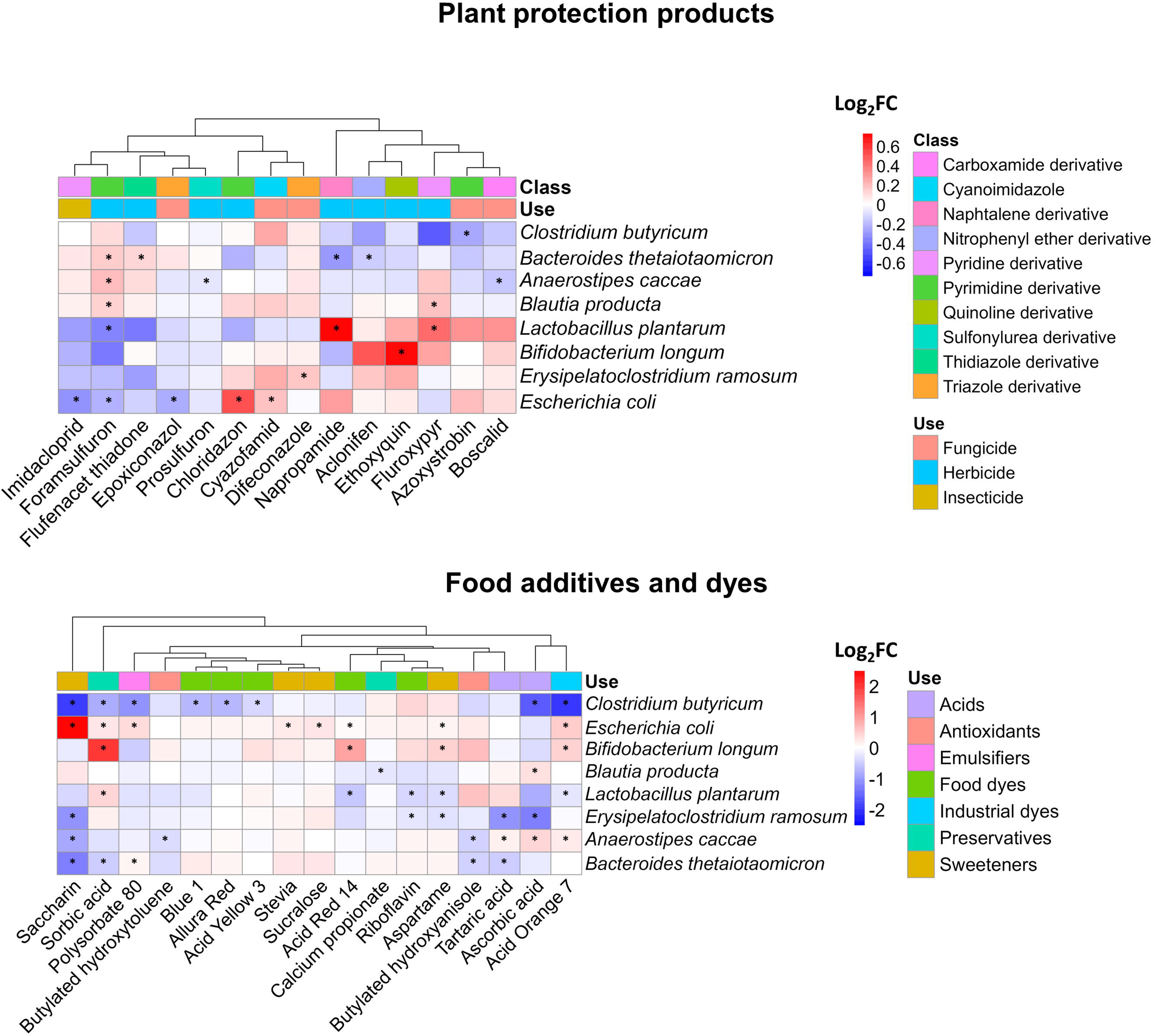
Impact of plant protection products, food additives, and dyes on the composition of SIHUMIx. The species abundance was measured by metaproteomics and is displayed as log2FC for (**A**) 14 plant protection products and (**B**) 17 food additives and dyes which affected at least the abundance of one species compared to the control. Statistically significant effects (*P_adj_* <0.05) are highlighted by asterisks.

Additionally, the range of species influenced by food additives and dyes exhibited greater variability than that observed for plant protection products, as illustrated in **Figure 2B**. For instance, only seven xenobiotics had a discernible impact on a single species, whereas the remaining ten exhibited effects on as many as five distinct species within the SIHUMIx community. The most pronounced alterations affecting a total of five different species were attributable to saccharin and acid orange 7. Among the affected species, *Clostridium butyricum* and *Escherichia coli* were the most prominently affected. Notably, the fold changes in species abundance ranged from -1.9 to a maximum of 2.5, suggesting both inhibitory and stimulatory effects of food additives and dyes on the composition of the SIHUMIx community.

### 3.2 Xenobiotic exposure led to a reduction of SCFAs

The production of SCFAs reflects the primary energy metabolism of the microbiota, offering valuable insights into the impact of environmental stressors on bacterial communities. Of the nine measured SCFAs, only isocaproate was below the detection limit. Among the 54 plant protection products tested, 10 demonstrated a significant effect on at least one of the eight analyzed metabolites (**Fig. 3A**). Interestingly, all of these plant protection products led to a reduction in metabolite concentrations, except for Benzotriazole, which resulted in an increase in the concentrations of the three major SCFAs (acetate, butyrate, and propionate), which together constituted up to 97% of the total SCFA measured (**Supplementary Figure S2**). We observed a distinct clustering pattern among affected SCFAs. Specifically, the three primary SCFAs were consistently clustered together. However, exceptions to this clustering occurred in the cases of acetamiprid and ethoxyquin, where less abundant SCFAs were affected and formed a distinct cluster. Among the SCFAs, butyrate was the most frequently affected, observed in seven instances involving plant protection products, while ethoxyquin affected the highest number of SCFAs, affecting six out of eight in total.

**Figure. 3:**
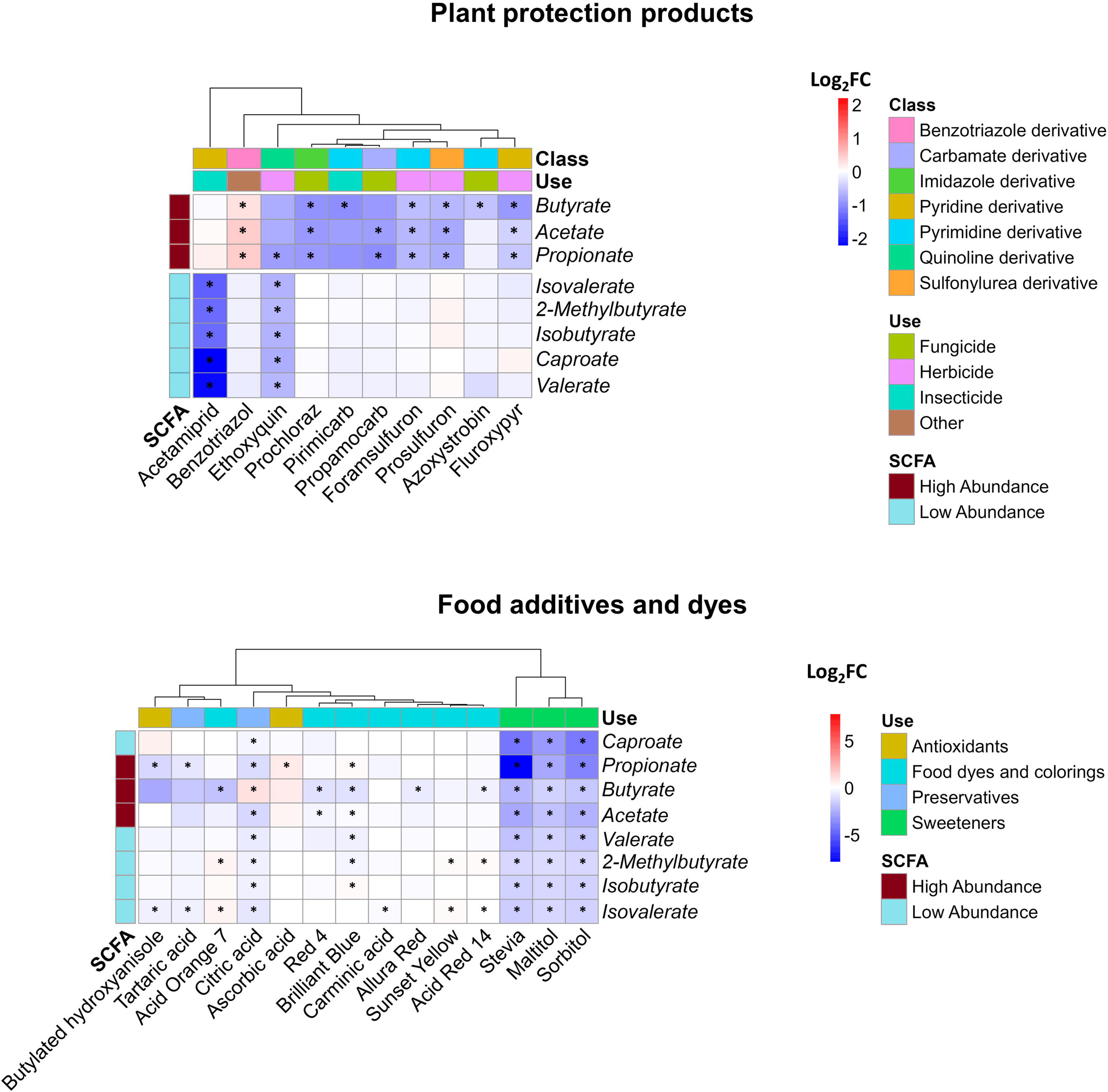
Impact of xenobiotics on SCFA concentration. Log2FC of the 8 detected SCFAs following 24h of exposure of SIHUMIx to (**A**) plant protection products and (**B**) food additives and dyes. Statistically significant effects (*P_adj_* <0.05) are indicated by asterisks.

No discernible correlations emerged among the plant protection product subgroups (**Fig. 3A**), namely insecticides, herbicides, and fungicides, as well as their effects on metabolites. This absence of correlation is also evident at the chemical structural level. Plant protection products, derived from diverse groups of chemicals including foramsulfuron, prosulfuron, prochloraz, primicarb, and propamocarb, influence the major SCFAs: acetate, propionate, and butyrate. In contrast to the previously mentioned cases, acetamiprid significantly reduced the concentration of the less abundant SCFAs without affecting the three major SCFAs. As an outlier, ethoxyquin decreased the levels of all measured metabolites, demonstrating a significant impact on minor SCFAs and propionate.

Concerning food additives, significant alterations were induced by 14 of the 36 tested compounds in at least one of the SCFAs, as depicted in **Figure 3B**. In contrast to the effects observed after exposure to plant protection products, the effects of food additives exhibit a mixture of results with a higher magnitude. For instance, propionate levels showed a remarkable decrease, reaching a fold change as low as -7.8 due to stevia, in contrast to pesticides, where the lowest value of -2.2 observed after exposure to acetamiprid. Isovalerate is the most frequently affected SCFA (10 out of 14 compounds), and the xenobiotics that cause the most significant effects are artificial sweeteners and citric acid, which altered all SCFAs.

Among the chemicals studied, ascorbic acid and sunset yellow exclusively led to significant increases in SCFA concentrations, whereas acid orange 7, citric acid, blue brilliant 1, and acid red 14 caused both elevations and reductions in the measured metabolite concentrations. Among the remaining xenobiotics, a consistent decrease in SCFA levels was observed. When categorizing the effects on SCFAs, it becomes evident that unlike plant protection products, the specific type of SCFA affected by these chemicals was not readily distinguishable. Instead, the use of chemicals to which the treatment belongs emerged as an influential factor. This is supported by the similar impacts of chemicals within the artificial sweeteners and food and industrial dye groups on the SCFA profile produced by SIHUMIx. We observed a distinct clustering of two xenobiotic groups based on their effects on SCFA production, suggesting that the chemicals within these groups might share a similar mode of action. The second group, which consisted of only sweeteners (stevia, maltitol, and sorbitol), consistently reduced all SCFAs and affected their relative abundances, with particular effects on propionate, acetate, butyrate, and caproate (**Supplementary Figure S3**).

### 3.3 Effects on metabolic pathways

To gain a more comprehensive understanding of community-level functionality, we performed a metaproteomic analysis. Following an examination of the treatments using the pathway selection algorithm outlined in the methods section, we discovered that 12 plant protection products and 22 food additives had a notable effect on at least one metabolic pathway. In total, 59 metabolic pathways exhibited significant alterations after exposure to plant protection products and food additives.

None of the 12 plant protection products that influenced SIHUMIx metabolism (**Supplementary Table S5**) belonged to the insecticide category. Of the 34 affected pathways affected only after exposure with plant protection products, 23 were affected by a single treatment, while the remaining 11 were affected by two treatments. Notably, propyzamide, prochloraz, and ethoxyquin had the most significant effects on these pathways (**Supplementary Figure S4**). To elaborate, propyzamide affected nine pathways, whereas prochloraz and ethoxyquin each affected eight pathways. It is important to highlight that prochloraz and propyzamide, along with flufenacet ESA and benzotriazole, increased the intensity of proteins in the affected pathways. In contrast, ethoxyquin and other significant treatments had a negative effect on the pathways.

We observed significant changes (*P_adj_* <0.05) in pathways related to xenobiotic response, regulation of gene expression, and antioxidant defense, including selenocompound metabolism, drug metabolism (propyzamide, log_2_FC = 0.5), and bacterial secretion system pathways (azoxystrobin, log_2_FC = -0.2). Additionally, we noted alterations in the basic energy and structural pathways, such as carbon metabolism (aclonifen, log_2_FC = -0.1), nitrogen metabolism (propyzamide, log_2_FC = 0.2), and pyruvate metabolism (ethoxyquin, log_2_FC = -0.1). The pentose phosphate pathway (prochloraz, log_2_FC = 0.1) and the metabolism of cysteine and methionine (fluroxypyr, log_2_FC = -0.1) and valine, leucine, and isoleucine (ethoxiquin, log_2_FC = -0.1) were also significantly affected.

We observed instances where the relationship between energy pathways and the production of SCFAs is evident. An example is ethoxyquin, where pathways such as starch and sucrose metabolism and pyruvate metabolism (log_2_FC =-0.1, *P_adj_* =0.028) exhibited reduced activity compared to the control group after exposure (**Fig. 4A.1**). This treatment also had a negative effect on SCFA production (**Fig. 4A.2**).

**Figure 4:**
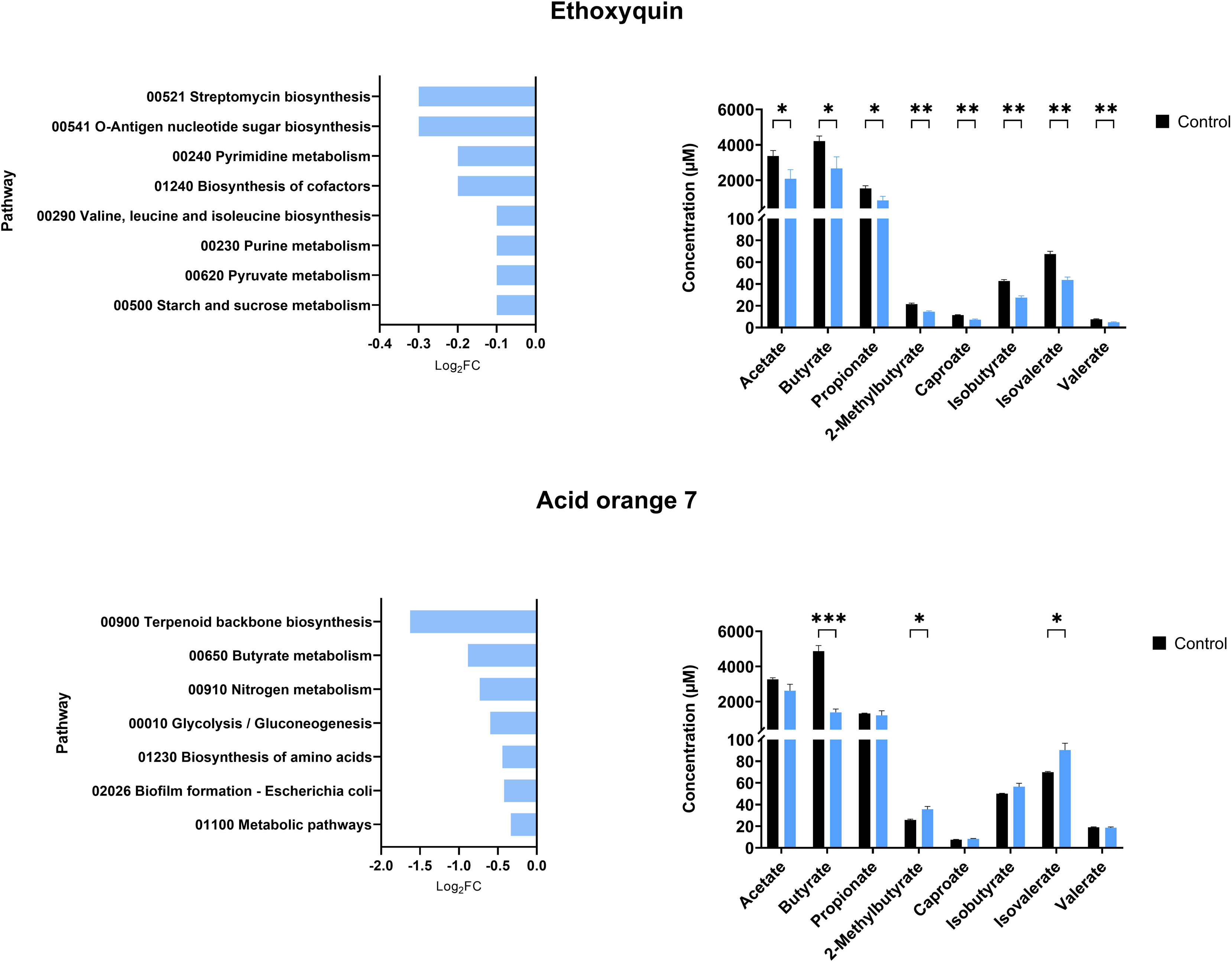
Impact of xenobiotics on SIHUMIx pathways and their relationship with SCFA production. Effects of **(A)** ethoxyquin and **(B)** acid orange 7 exposure on SIHUMIx significantly **(1)** affected pathways and their association with changes in **(2)** short-chain fatty acid (SCFA) concentrations. Following exposure to the plant protection product, we observed a reduction in energy pathways, notably the metabolism of pyruvate, a key precursor for SCFAs. Acid orange 7, on the other hand, led to a decline in butyrate metabolism, resulting in a significant decrease in butyrate concentration levels in the supernatant. Statistically significant effects (*P_adj_* <0.05) are indicated by asterisks.

Similar to the findings from the SCFA analysis, we observed that food additives and dyes had a more significant impact on the scale and diversity of metabolic pathway effects, whereas exposure to plant protection products exhibited a narrow range with a minimum log_2_FC value of -0.4 and a maximum of 0.6 (**Supplementary Table S5**). In contrast, exposure to food additives and dyes resulted in an approximately ten-fold effect, with minimum values of -3.2 and a maximum of 5.7 (**Supplementary Table S6**). Among the 36 treatments, 22 induced significant alterations in 61 metabolic pathways constituting 61% of all the food additive and dye treatments. Within this group, seven treatments (isomalt, carrageenan, polydimethylsiloxane, aspartame, riboflavin, polysorbate 80, and stevia) exclusively increased the intensity of metabolic pathways. Conversely, allura red and ascorbic acid alone led to a reduction in proteins associated with metabolic pathways. The remaining 13 treatments produced mixed outcomes, revealing both increases and decreases in the pathway activity.

A comprehensive overview of the treatments and their impacts on various pathways is presented in **Supplementary Figure S5**. Notable highlights include the most significantly affected pathway, which experienced a decrease due to 11 different treatments: resistance to cationic antimicrobial peptides (cAMP). These peptides are produced by the host to defend against bacterial infection. Additionally, several other pathways were affected by multiple treatments, including the cell cycle, 2-oxocarboxylic acid metabolism, histidine metabolism, thiamine metabolism, and the pantothenate and CoA biosynthesis pathways. Among the treatments, saccharine caused the greatest disturbances, significantly affecting 15 pathways. These included lipoic acid metabolism (log_2_FC = 5.7), sulfur relay system (log_2_FC = 3.8), and terpenoid backbone biosynthesis (log_2_FC = -2.2).

We also observed instances where the outcomes of affected pathways were correlated with the results of short-chain fatty acids (SCFAs). For example, ethoxyquin, a treatment that led to a decrease in the concentration of all SCFAs (**Fig. 4A.2**), exhibited a reduction in the starch and pyruvate metabolism pathways (log_2_FC = -0.1, *P_adj_* = 0.028), as well as in the biosynthesis of branched amino acids (log_2_FC = -0.1, *P_adj_* = 0.046) **(Fig. A.1)**, the latter serving as the foundation for the production of branched SCFAs. Similarly, acid orange 7 showed a decline in the metabolic pathway associated with butyrate production (**Fig. 4B.1**) (log_2_FC = -0.9, *P_adj_* = 0.017), aligning with the decrease observed in butyrate measurements (**Fig. 4B.2**).

A correlation analysis was performed to investigate the potential associations between the relative abundance of SIHUMIx members and the metabolic pathways influenced by xenobiotic exposure. Only correlations with a Pearson’s index higher than 0.6 and lower than -0.6, with *P_adj_* <0.05, were considered significant. In the context of plant protection products (**Fig. 5A**), two metabolic pathways were positively correlated with the three species: the sulfur relay system pathway exhibited a correlation with *E. ramosum* (r = 0.696, *P_adj_* <0.001) and *E. coli* (r = 0.619, *P_adj_* <0.001), while the terpenoid backbone biosynthesis pathway showed a correlation with *C. butyricum* (r = 0.677, *P_adj_* <0.001). Conversely, *B. longum* showed a negative correlation with drug metabolism, other enzymes (r = -0.652, *P_adj_* <0.001), and oxidative phosphorylation (r = -0.746, *P_adj_* <0.001). The latter pathway was also found to be negatively linked with *L. plantarum* (r = -0.633, *P_adj_* <0.001).

**Figure 5:**
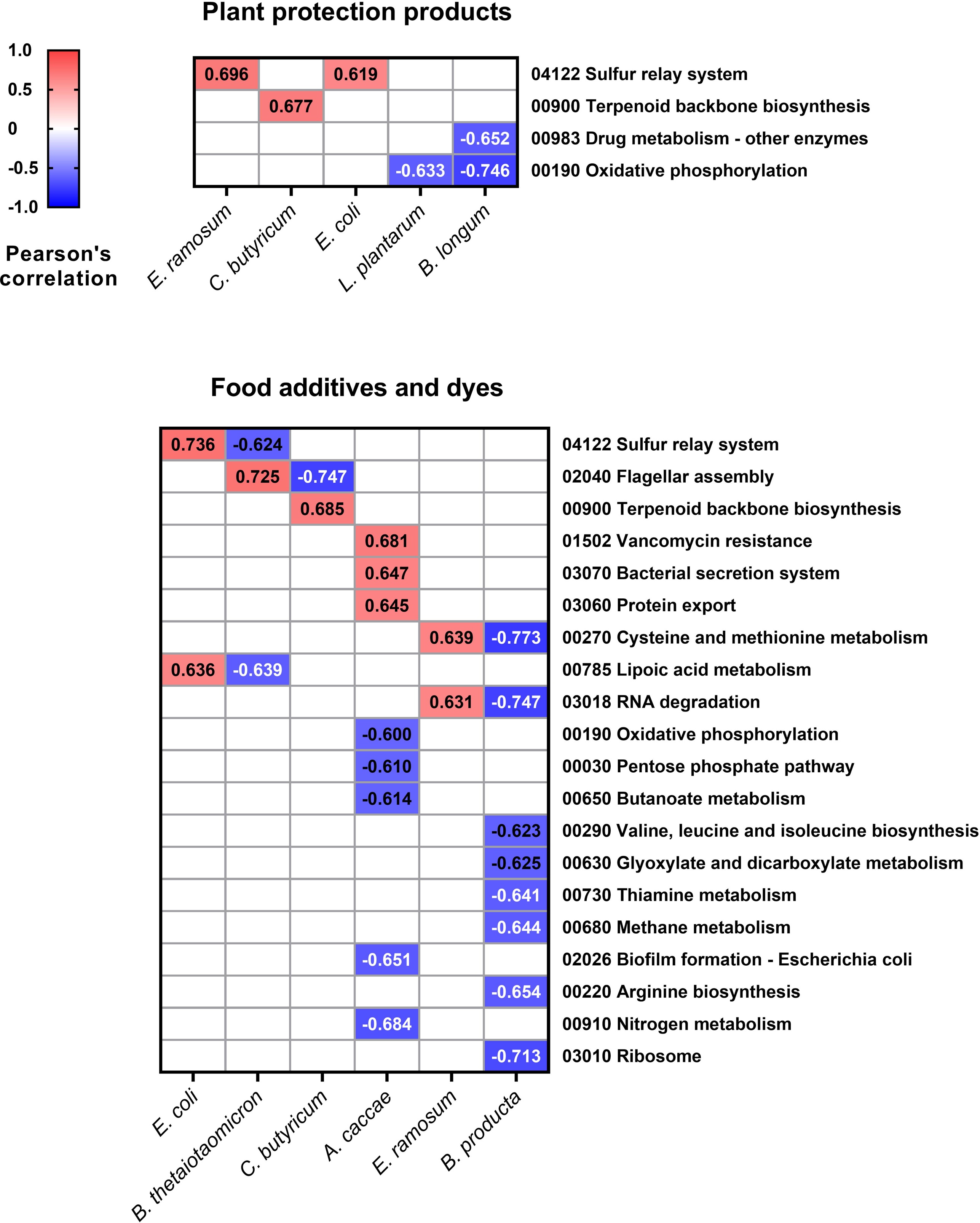
Pearson’s correlation values between relative species abundance and affected pathways. Values for the relative abundance of SIHUMIx species and pathways lower than -0.6 and higher than 0.6 and *P_adj_* <0.05 after exposure to (**A**) plant protection products and (**B**) food additives and dyes.

Regarding food additives, positive correlations were identified between nine metabolic pathways and five species of SIHUMIx (**Fig. 5B**). In particular, there was a positive correlation between *A. caccae* and vancomycin resistance (r = 0.681, *P_adj_* <0.001), along with pathways associated with the efflux of substances from the cell: bacterial secretion system (r = 0.647, *P_adj_* <0.001) and protein export (r = 0.645, *P_adj_* <0.001). Conversely, 16 metabolic pathways were negatively correlated with four SIHUMIx species. *B. producta* demonstrated eight negative correlations, with pathways linked to the synthesis of amino acids such as valine, leucine, and isoleucine (r = -0.623, *P_adj_* <0.001) as well as arginine biosynthesis (r = -0.654, *P_adj_* <0.001) standing out prominently.

## 4 Discussion

In this study, we employed metaproteomics and metabolomics to investigate the effects of 90 xenobiotics on Simplified Human Gut Microbiota (SIHUMIx). We exposed stable SIHUMIx aliquots to 54 plant protection products and 36 food additives and dyes at environmentally relevant concentrations in 96-well-deep plates. Through metaproteomics and metabolomics, we examined the changes in species abundance, metabolic pathway intensity, and SCFA production (**Figure 1**). Our primary objective was to explore the impact of these chemicals on bacterial microbiota models, a less common aspect of risk assessment, and to show their potential significance for human microbiota and host health.

We evaluated the effects of two groups of chemicals that encompass a wide array of structures and functional differences. Despite variations in their levels of exposure and toxicity, both groups exhibit discernible effects on human health (Juraske et al., 2009; Cao et al., 2020). To quantify the extent of exposure to these xenobiotics, we calculated the concentration adjusted for the number of bacterial cells in our model relative to the total cells in the daily fecal output. The plant protection products were administered to SIHUMIx at a median concentration of 0.01 mg/mL (**Supplementary Table S2**), whereas the food additives and dyes were introduced into the culture at a median of 1.03 mg/mL (**Supplementary Table S3**). These concentration ranges differ significantly, which potentially accounts for the disparities in the extent and magnitude of these effects between the two groups. Nevertheless, these variations accurately mirror the levels of exposure in the environment and produce.

### 4.1 Xenobiotics affected SIHUMIx structure

Out of the 14 plant protection products that had an impact on at least one SIHUMIx species (**Fig. 2A**), eight have not yet been tested in any animal microbiota model. These products include fluroxypyr, aclonifen, chloridazon, cyazofamid, napropamide, prosulfuron, flufenactet ESA, and foramsulfuron; making this the first study on the influence of these xenobiotics on a microbiota model. On the other hand, the remaining six chemicals have been examined in microbiota models, which differ significantly from human models. The reported effects in the current literature showed changes in abundance at phyla level caused by azoxystrobin (Meng et al., 2022), ethoxyquin (Wei et al., 2023), boscalid (Dong et al., 2023), and imidacloprid (Yang et al., 2020), as well difeconazole (Bao et al., 2022), and epoxiconazole (Xu et al., 2014). Our study facilitated the identification of alterations in the community structure at the species level. These modifications in microbiota diversity may indicate a disturbance in the host’s well-being, leading to a heightened risk of autoimmune and inflammatory diseases (Blaser, 2017); while also creating opportunities for infection by opportunistic bacteria (Lichtman et al., 2016). Therefore, the changes observed after exposure to agroindustrial xenobiotics could improve our understanding of the direct toxicity of chemicals beyond genus level.

The utilization of artificial sweeteners has gained increasing popularity in recent years as a substitute for conventional sweetening agents, and so has interest in their effects on the microbiome. Investigations have identified potential associations between the consumption of these substances and disruptions in gastrointestinal health (Shil et al., 2020) as well as their limited efficacy in facilitating weight loss (Brown et al., 2010). *E. coli* showed significant increases in its relative abundance after being exposed to Non-caloric artificial sweeteners (NAS) such as saccharin (log_2_FC = 2.5), stevia (log_2_FC = 0.2), sucralose (log_2_FC = 0.3), and aspartame (log_2_FC = 0.5). This result agrees with previous research, which demonstrated a direct impact of this class of artificial sweeteners on *E. coli* (Shil and Chichger, 2021), stimulating its proliferation and eliciting heightened expression of enzymes linked to energy metabolism, fatty acid biosynthesis, and amino acid metabolism (Mahmud et al., 2020).

Some species belonging to the phylum Firmicutes exhibited a reduction in their relative abundance following exposure to various xenobiotics. *L. plantarum* showed decreased relative abundances after being exposed to the dye acid orange 7 (log_2_FC = 0.3), riboflavin (log_2_FC = -0.4), acid red 14 (log_2_FC = -0.5), and the sweetener aspartame (log_2_FC = -0.3); while *C. butyricum*, was affected by eight treatments, some examples being saccharin (log_2_FC = -1.9), acid orange 7 (log_2_FC = -1.9), and ascorbic acid (log_2_FC = -1.6) (**Fig. 2B**). These species play pivotal roles in maintaining intestinal health and microbiota stability, and their reduction in abundance could pose a risk to balance and gut health. *C. butyricum*, is known as a butyrate producer (He et al., 2005), has the capacity to alleviate inflammation, maintain intestinal homeostasis (Pan et al., 2019; Ma et al., 2022), modulate immune responses (Chen et al., 2019), and promote the growth of beneficial bacteria from genera such as *Bifidobacterium* and *Lactobacillus*. *L. plantarum* is a probiotic species that contributes to microbiota rebalancing by mitigating the overgrowth of bacteria belonging to the Enterobacteriaceae phylum (Linninge et al., 2019), reinforcing the intestinal barrier (Zhao et al., 2020), and counteracting the effects of other toxic compounds on the microbiota, such as heavy metals (Yu et al., 2021).

### 4.2 Changes in SCFAs concentrations were observed after xenobiotics exposure

Short-chain fatty acids, which are the primary byproducts of dietary fiber, proteins, and peptides, are recognized as immunomodulatory agents and supportive elements for host well-being. Notably, the gut microbiota predominantly produces acetate, propionate, and butyrate as major products (Blaak et al., 2020).

Various responses to SCFA production were observed across different plant protection products (**Fig. 3A**). We observed a reduction (log_2_FC <-0.5) in acetate, butyrate, and propionate after exposure to foramsulfuron which reduced the abundance of *L. plantarum* (log_2_FC = -0.3) and in butyrate after exposure to azoxystrobin affecting *C. butyricum* (log_2_FC = -0.2). Alterations in bacterial composition, specifically reduction in the abundance of SCFA-producing species, could be the cause of the observed reduction in concentrations (Kusumo et al., 2019; Hsiao et al., 2021). Conversely, in the cases of ethoxyquin and fluroxypyr, there was no significant decrease of the abundance of SCFA-producing species but in contrast, we observed a reduction of the intensity of proteins involved in the energy metabolism (log_2_FC = -0.2) and amino acid metabolism (log_2_FC = -0.1), which are key pathways for SCFA production (Morrison and Preston, 2016). Acetamiprid, pirimicarb, and propamocarb decreased SCFA concentrations compared with the control, but no effects on the structure and metabolic pathways were observed. Further analysis is needed to confirm whether this is due to the loss of the producing species or metabolic alterations.

The adverse effects of azo dyes on the microbiota, altering its structure and reducing SCFA production, have been described previously (Polic, 2018; Wu et al., 2021). These studies have established that azo dyes induce changes in bacterial composition, diminish SCFA production, and produce potentially carcinogenic compounds following degradation by azoreductases. In our study, these effects were readily observable in treatments that fell within the primary cluster of dyes (**Fig. 3B**), primarily influencing butyrate, 2-methylbutyrate, and isovalerate levels. Notably, among this group, only acid red 14 and brilliant blue exhibited additional impacts on the relative abundances of SCFA-producing species of SIHUMIx, by causing a decrease in *C. butyricum* (log_2_FC = -0.7) and *L. plantarum* (log_2_FC = -0.5). For the industrial dye acid orange 7, which also demonstrated a decline in SCFA-producing species (*C. butyricum*, log_2_FC = -1.9), we observed a negative effect on the butanoate metabolism pathway (log_2_FC = -0.9). Among the proteins detected in this pathway in comparison to the control group, the enzyme acetyl-CoA transferase was undetectable, and there was a discernible reduction in the activity of enzymes, such as butyryl-CoA dehydrogenase, acetyl-CoA C-acetyltransferase, and butyrate kinase.

A notorious cluster effect affecting all SCFAs was evident when SIHUMIx was exposed to sweeteners, leading to a decrease in the concentration of all detected metabolites (**Fig. 3B**). SCFA production stems from fermentation of resistant, non-digestible starch (Flint et al., 2012). Substituting these starches with rapidly absorbed sources, such as artificial sweeteners, could potentially have a detrimental impact on SCFA production (Scott et al., 2008; Farup et al., 2019). In our study, we did not remove complex carbohydrate sources; however, it is possible that SIHUMIx shifted their primary carbohydrate source towards artificial sweeteners, resulting in an overall reduction of all SCFAs.

### 4.3 Xenobiotic exposure influenced SIHUMIx on functional level

The metaproteomic analysis has unveiled significant alterations and correlations between pathways and the relative abundance of certain species within the SIHUMIx microbiome. Notably, *E. coli, C. butyricum,* and *E. ramosum* exhibited positive correlations with pathways that can indirectly intersect within the broader context of cellular metabolism (**Fig. 5A**). For instance, the sulfur relay system may provide the essential sulfur atoms needed for synthesizing specific cofactors and coenzymes crucial in terpenoid biosynthesis (Mihara and Esaki, 2002). Furthermore, terpenoids themselves can undergo modification through enzymatic reactions that may necessitate sulfur-containing cofactors (Kotera et al., 2010). Conversely, the reduction in lactic acid bacteria such as *L. plantarum* and *B. longum* (**Fig. 5A**) could imply a detriment to energy production, as well as compromised detoxification, resistance, and biotransformation of xenobiotics as mentioned earlier. This is evident from their negative correlation with drug metabolism and oxidative phosphorylation pathways.

Regarding the magnitude of effects of plant protection products, prochloraz, and ethoxyquin, with eight pathways affected, and propyzamide, with nine, were the treatments that impacted most pathways (**Supplementary Figure S4**). The effects of these chemicals on microbiota have been studied in other models, such as mice (Wang et al., 2021), cow (Wei et al., 2023), and zebrafish (Sanmarco et al., 2022). After exposure to propyzamide and prochloraz, there were notable increases in energy-related pathways (log_2_FC >0.1), such as fructose and mannose metabolism, nitrogen metabolism, and pentose phosphate pathways. These increases may indicate a higher energy demand by the cell to cope with the effects of these two pesticides (Aertsen and Michiels, 2004). Conversely, after exposure to ethoxyquin we observed a decrease in pathways associated with energy metabolism (starch, sucrose and pyruvate; log_2_FC = -0.1) and DNA synthesis, affecting the metabolism of purines (log_2_FC = -0.1) and pyrimidines (log_2_FC = -0.2). However, it is crucial to emphasize that further research is required to fully understand the effects of these chemicals on human and human-related microbiota models, as well as their potential risks to host health.

In regard to the impact of food additives and dyes, it has been observed that they have a more pronounced effect on the scale and diversity of metabolic pathways when compared to plant protection products (**Supplementary Figure S5**). Notably, the resistance to the cationic antimicrobial peptides (cAMP) pathway is significantly affected, experiencing a reduction due to 11 different treatments. This decrease in resistance to cAMP suggests a higher vulnerability of microbiota species to host defenses, providing an opportunity for opportunistic bacteria to colonize and establish infections, potentially posing a risk to the host’s health (Band and Weiss, 2014).

The *B. producta* strain exhibited a significant negative correlation with nine metabolic pathways (**Fig. 5B**), with a particular emphasis on pathways associated with protein synthesis, including cysteine and methionine metabolism, valine, leucine, and isoleucine biosynthesis, arginine biosynthesis, and ribosomal activity. This can occur through the activation of xenobiotic stress response pathways, modification of oxidative phosphorylation function, and the redirection of resources toward xenobiotic metabolism (Richardson et al., 2013; Chen et al., 2019). Similarly, regarding xenobiotic metabolism, *A. caccae* showed positive correlation values with pathways involved in the extrusion of compounds outside the bacterial cell (**Fig. 5B**). These stress response pathways, which are over-regulated, play a crucial role in the bacterial response to xenobiotics (Riesbeck et al., 2022), enabling bacteria to export effector proteins, promote bacterial attachment, and adapt to changing environments.

## 5 Conclusions

The use of an *in vitro* bioreactor system that enabled continuous SIHUMIx production in combination with a high-throughput model using 96-well deep plates allowed the detection of specific changes in key aspects of a human microbiota model after exposure to a wide range of xenobiotics. Alterations in species-level abundances, changes in metabolic pathways, and SCFAs concentrations were noticeable, even at concentrations as low as 0.00175 mg/mL (flufenacet ESA, **Supplementary Table S2**) using metabolomics and metaproteomics, the latter providing both structural and functional information surpassing the predictive limitations of techniques such as metagenomics.

Our approach is not free of limitations. First, the complexity of our microbial community, although it represents the functional core of the human microbiome (Becker et al., 2011), is low and does not reflect the microbial diversity found in human fecal samples (Li et al., 2019b) or in highly complex synthetic communities (Perez et al., 2021). However, these may exhibit greater variability, depending on the nature of the donor or cultivation system (Johnson et al., 2020). Second, our current implementation measures only the direct effects of the compounds on the microbiome. It does not consider the impact of the host on the microbiome or the effects of metabolites produced by the host.

The wide array of results obtained demonstrates the versatility of the method when it comes to exposing cultivable communities to a large number of chemicals, or the potential to expose a single chemical to several concentrations in a short timeframe. The combination of this high-throughput method with the stability of SIHUMIx provides a suitable choice for risk assessment studies where there is an urgent need to elucidate the chemical’s impact on the microbial community and the role of microbiota (Ampatzoglou et al., 2022).

In this study, we used metaproteomics and metabolomics techniques to assess the impact of 90 xenobiotics on SIHUMIx, employing a combination of *in vivo* cultivation strategies. This comprehensive analysis covered 54 plant protection products and 36 food additives and dyes, effectively replicating the environmentally relevant concentrations. Our study revealed notable effects on the microbial abundance of SIHUMIx and its metabolic pathways as well as SCFAs.

These effects may indicate a compromise in the functionality of the microbiota, potentially jeopardizing its symbiotic relationship with host organism. Substantial variations in both the quantity and magnitude of these effects were observed between the two distinct groups of chemicals, underscoring the importance of accounting for exposure levels when comprehending their respective consequences. This investigation provides novel insights into the influence of xenobiotics on the human microbiota, with potential implications for autoimmune disorders, infections, and gastrointestinal health. Furthermore, the methodology employed in this study is proposed as a valuable tool for the assessment of chemical risks, providing a systematic means to evaluate the impact of xenobiotics on human microbiota. In turn, this contributes to a more comprehensive understanding of the potential ramifications of human health and well-being.

## 6 Author Contributions

**Conceptualization and Methodology,** V.C.-M., N.J., L.-F.F., and Q.F.; **Investigation,** V.C.-M., and L.-F.F.; **Data curation**, V.C.-M., S.-B.H., and M.N.H.; **Formal analysis,** V.C.-M. and M.N.H.; **Supervision**, N.J., U.R.-K., M.v.B., and Q.F.; **Visualization**,V.C.-M, S.-B.H, M.N.H.; **Writing—original draft**, V.C.-M., N.J., and M.v.B. ; **Writing-review and editing,** V.C.-M., L.-F.F., Q.F., S.- B.H., U.R.-K., M.N.H., M.v.B, and N.J.

## Acknowledgments

We want to thank Kathleen Eismann and Nicole Bock for their excellent technical assistance, and Kristian Jensen Pedersen, Cassandra Uthoff, Zheng Chen, Sebastian Gutsfeld, and Tamara Tal for their support and feedback.

## 9 Data Availability Statement

The mass spectrometry proteomics data have been deposited to the ProteomeXchange Consortium via the PRIDE partner repository (Perez-Riverol et al., 2022) with the dataset identifier PXD047410.

