## Supplementary material for "High-Throughput Screening of the Effects of 90 Xenobiotics on the Simplified Human Gut Microbiota Model (SIHUMIx): A Metaproteomic and Metabolomic Study": Figure S1

**Figure S1: Concentration of sulfasalazine over the course of seven days during the screening process.** Biotic samples comprised SIHUMIx exposed to 1µM sulfasalazine for 24 hours, while abiotic samples consisted of CIM media and 1µM sulfasalazine.

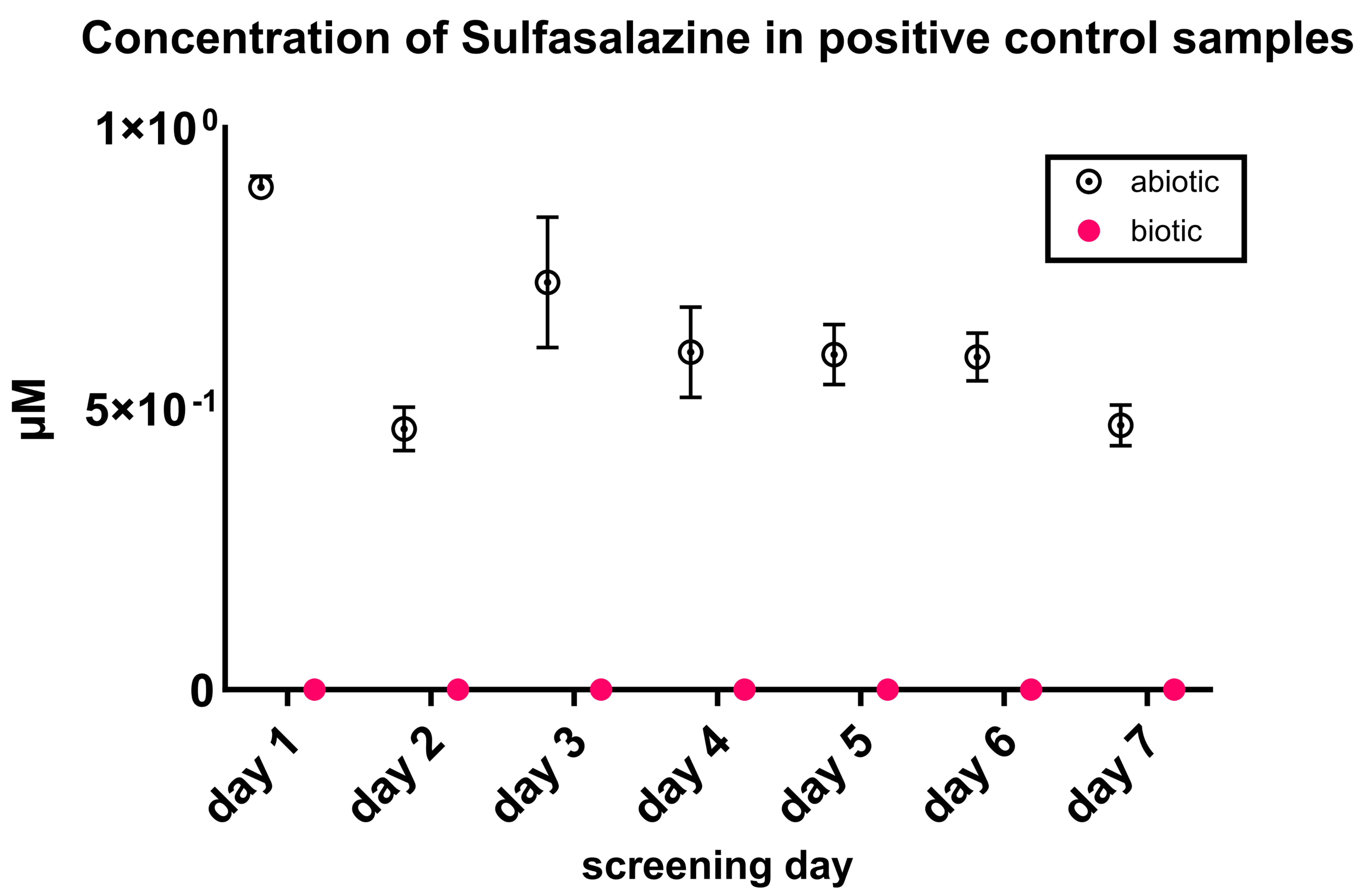
