## Supplementary material for "High-Throughput Screening of the Effects of 90 Xenobiotics on the Simplified Human Gut Microbiota Model (SIHUMIx): A Metaproteomic and Metabolomic Study": Figure S2

**Figure S2: Relative abundances of SCFAs after the exposure of SIHUMIx to Plant protection products:** High abundance (left side) and low abundance (right side) SCFAs after exposure to **(A)** acetamiprid and azoxystrobin, **(B)** fluroxypyr and foramsulfuron, **(C)** pirimicarb, prochloraz, propamocarb and prosulfuron, **(D)** benzotriazol, **(E)** ethoxyquin.

**A**

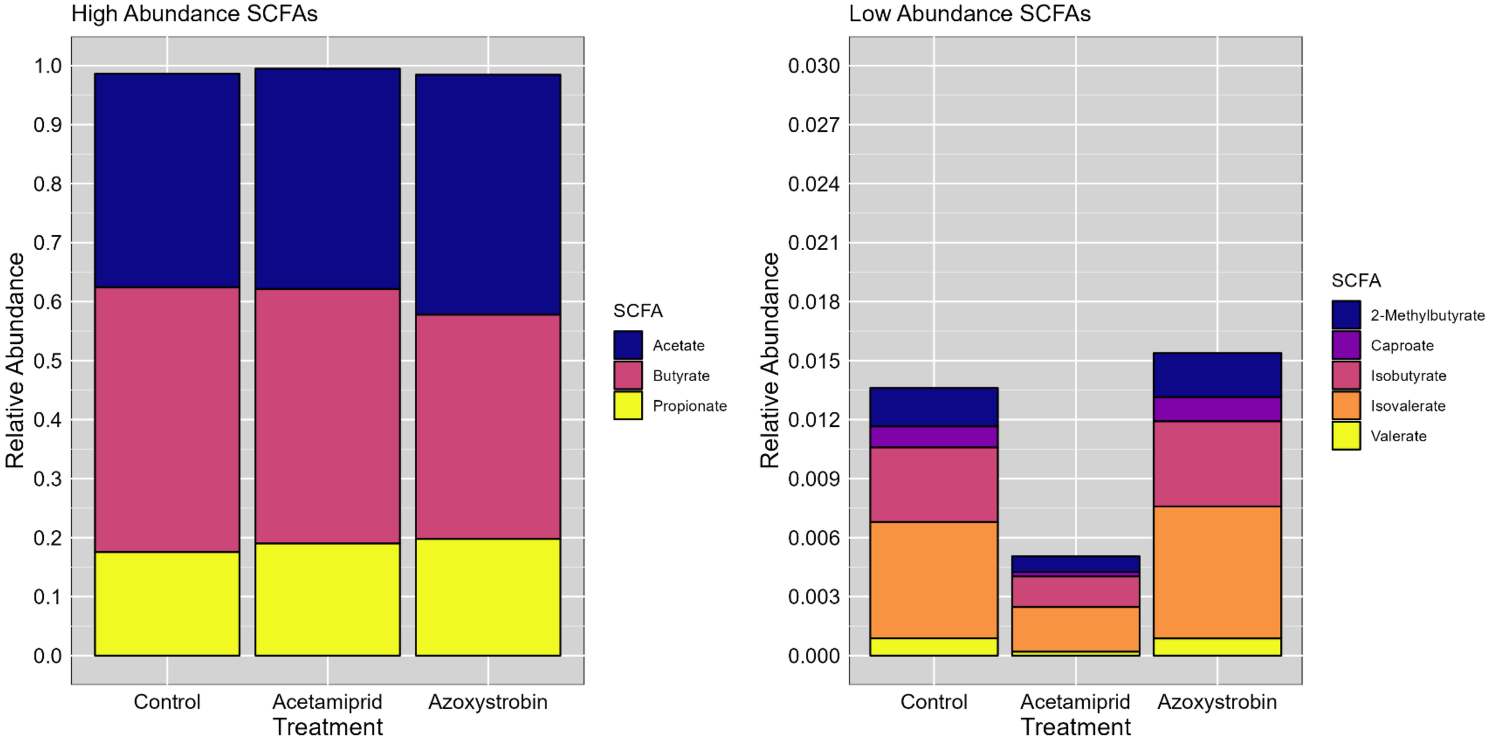


**B**

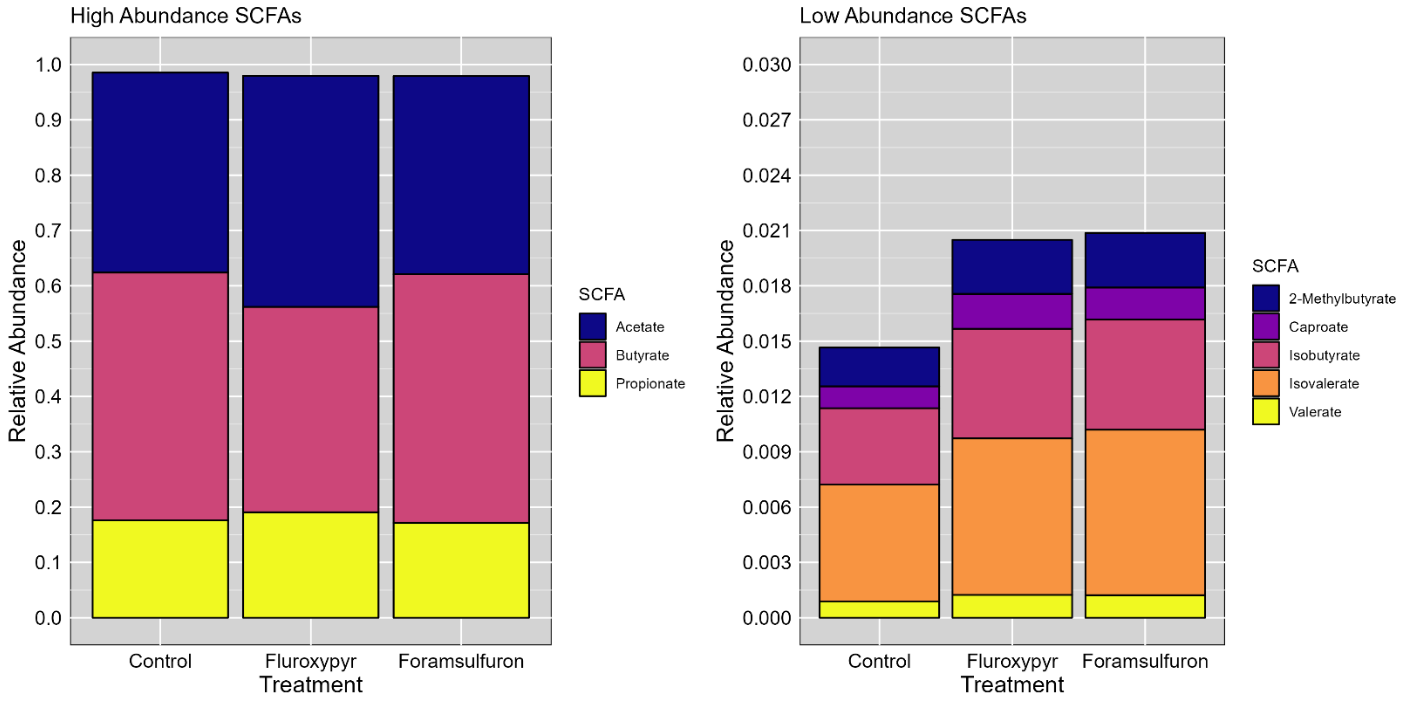

**C**

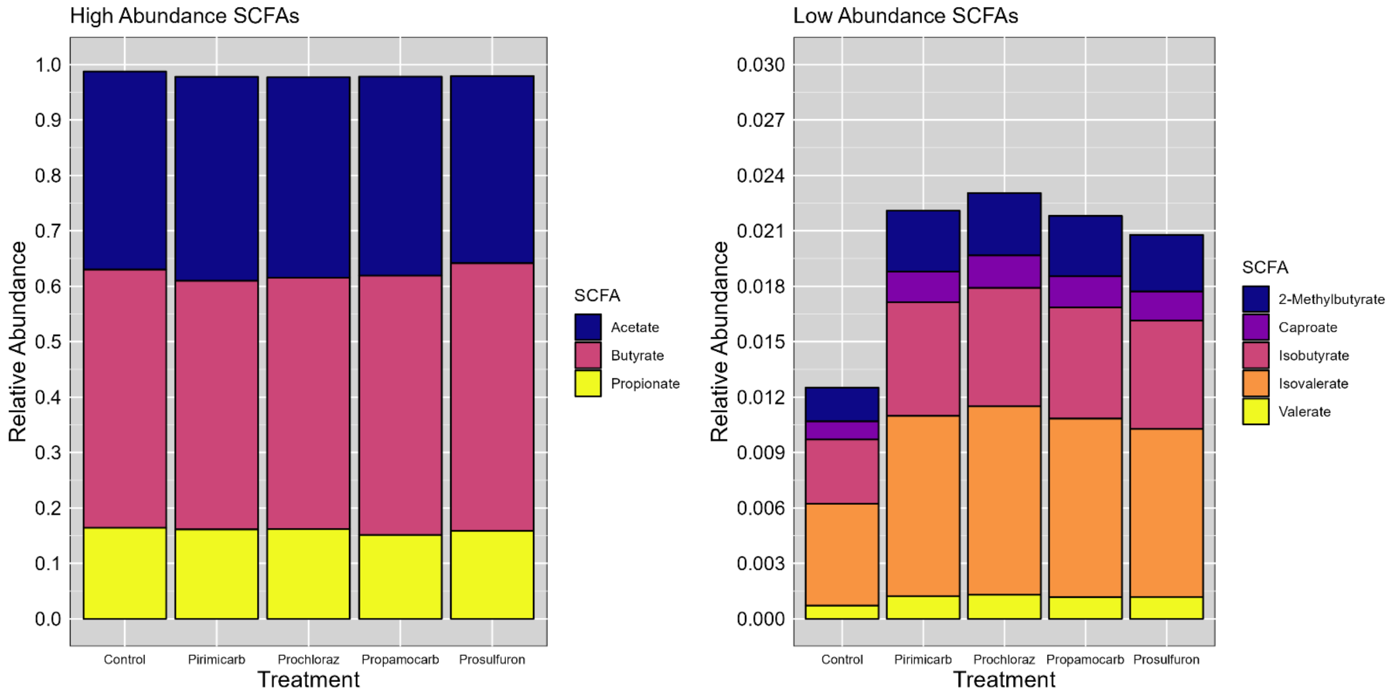
High Abundance SCFAs
Relative Abundance
1.0
0.9
0.8
0.7
0.6
0.5
0.4
0.3
0.2
0.1
0.0
Control
Pirimicarb
Prochloraz
Propamocarb
Prosulfuron
Treatment
SCFA
Acetate
Butyrate
Propionate
Low Abundance SCFAs
Relative Abundance
0.030
0.027
0.024
0.021
0.018
0.015
0.012
0.009
0.006
0.003
0.000
Control
Pirimicarb
Prochloraz
Propamocarb
Prosulfuron
Treatment
SCFA
2-Methylbutyrate
Caproate
Isobutyrate
Isovalerate
Valerate

**D**

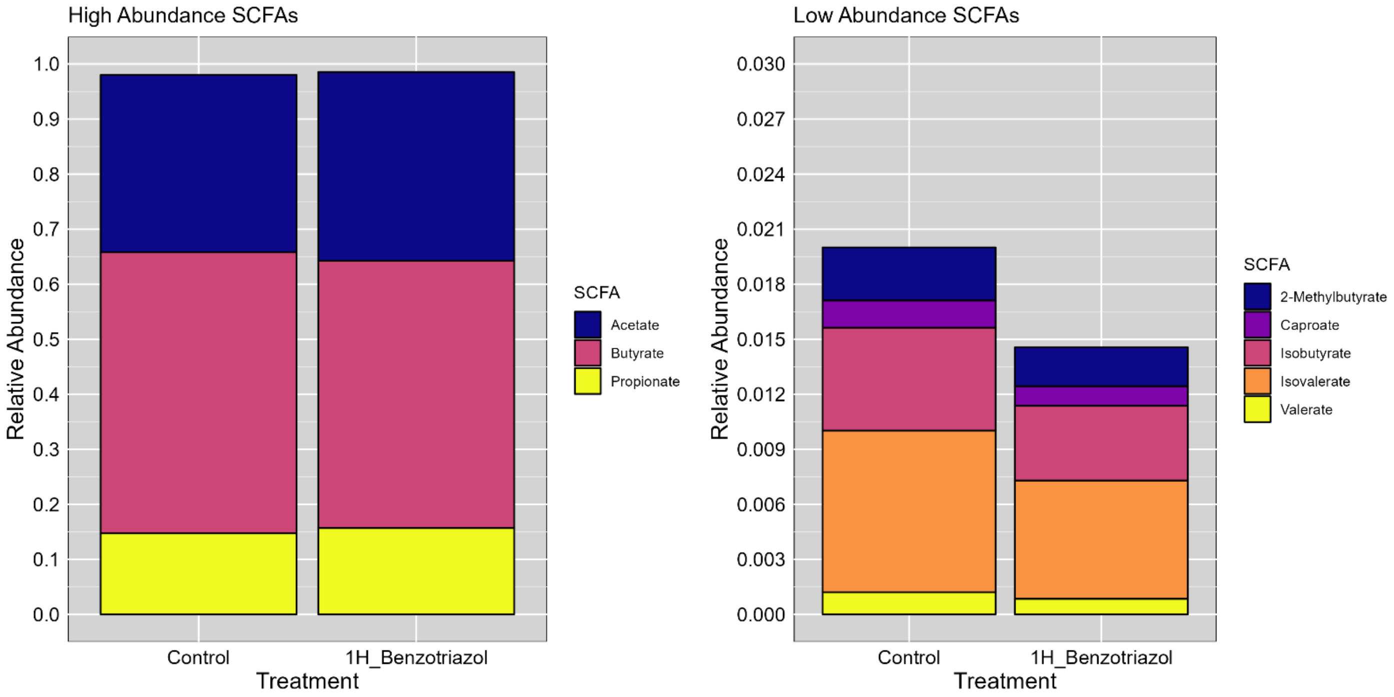
High Abundance SCFAs
Relative Abundance
1.0
0.9
0.8
0.7
0.6
0.5
0.4
0.3
0.2
0.1
0.0
Control
1H_Benzotriazol
Treatment
SCFA
Acetate
Butyrate
Propionate
Low Abundance SCFAs
Relative Abundance
0.030
0.027
0.024
0.021
0.018
0.015
0.012
0.009
0.006
0.003
0.000
Control
1H_Benzotriazol
Treatment
SCFA
2-Methylbutyrate
Caproate
Isobutyrate
Isovalerate
Valerate

E

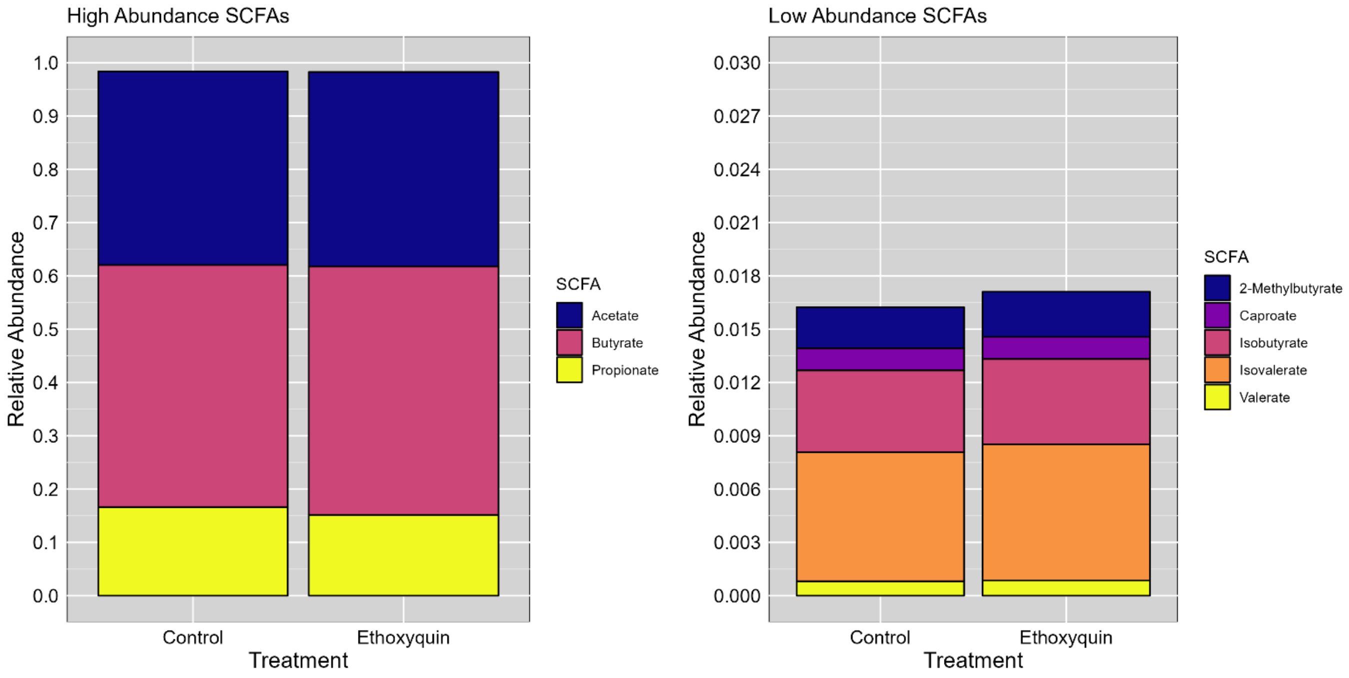

High Abundance SCFAs
1.0
0.9
0.8
0.7
0.6
0.5
0.4
0.3
0.2
0.1
0.0
Relative Abundance
Control
Ethoxyquin
Treatment
SCFA
Acetate
Butyrate
Propionate
Low Abundance SCFAs
0.030
0.027
0.024
0.021
0.018
0.015
0.012
0.009
0.006
0.003
0.000
Relative Abundance
Control
Ethoxyquin
Treatment
SCFA
2-Methylbutyrate
Caproate
Isobutyrate
Isovalerate
Valerate
