## Supplementary material for "High-Throughput Screening of the Effects of 90 Xenobiotics on the Simplified Human Gut Microbiota Model (SIHUMIx): A Metaproteomic and Metabolomic Study": Figure S3

**Figure S3: Relative abundances of SCFAs after the exposure of SIHUMIx to food additives and dyes:** High abundance (left side) and low abundance (right side) SCFAs after exposure to **(A)** butylated hydroxyanisole, sorbic acid and saccharine, **(B)** tartaric acid, citric acid and ascorbic acid, **(C)** cochineal, red 4, sunset yellow, acid red 14, acid orange 7, blue 1 and allura red, **(D)** maltitol, sorbitol and stevia.

**A**

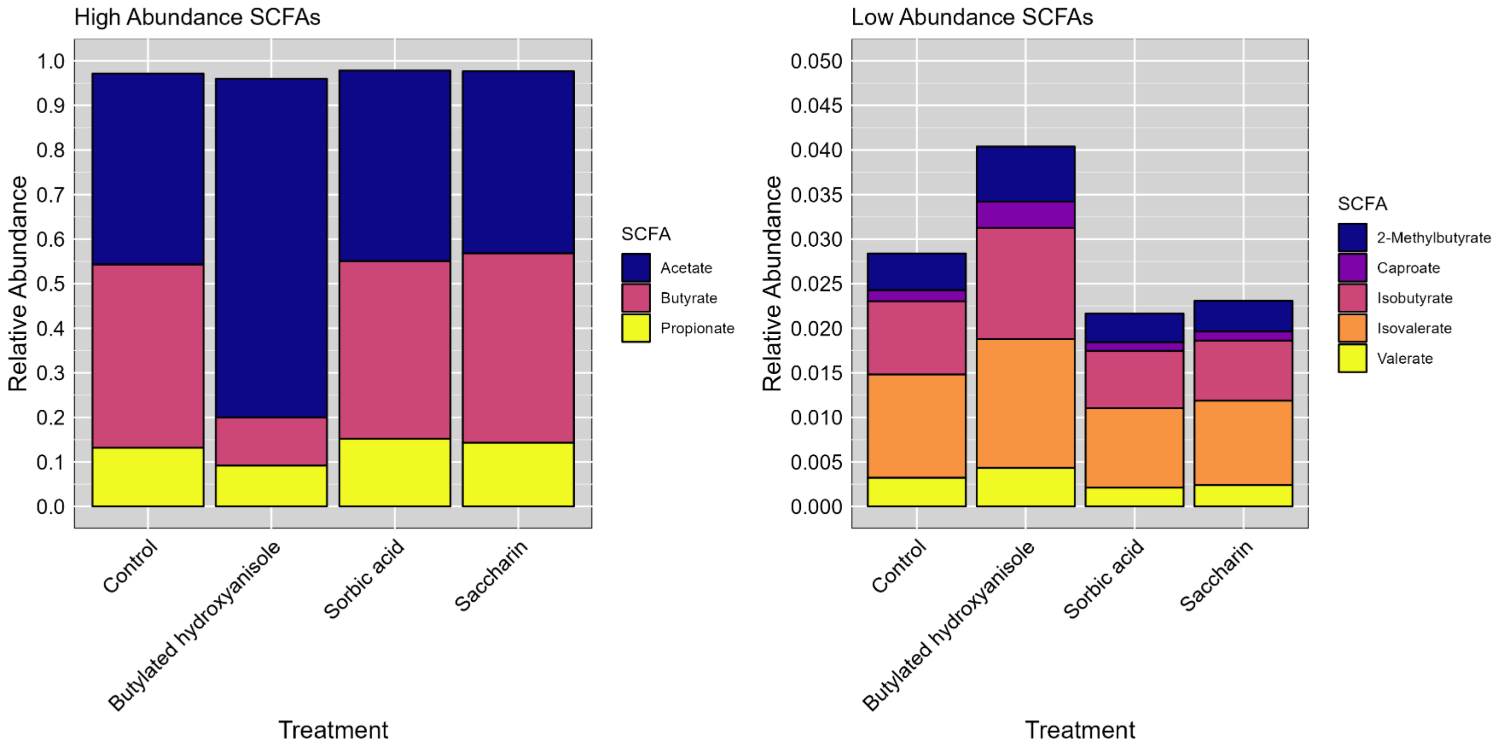


**B**

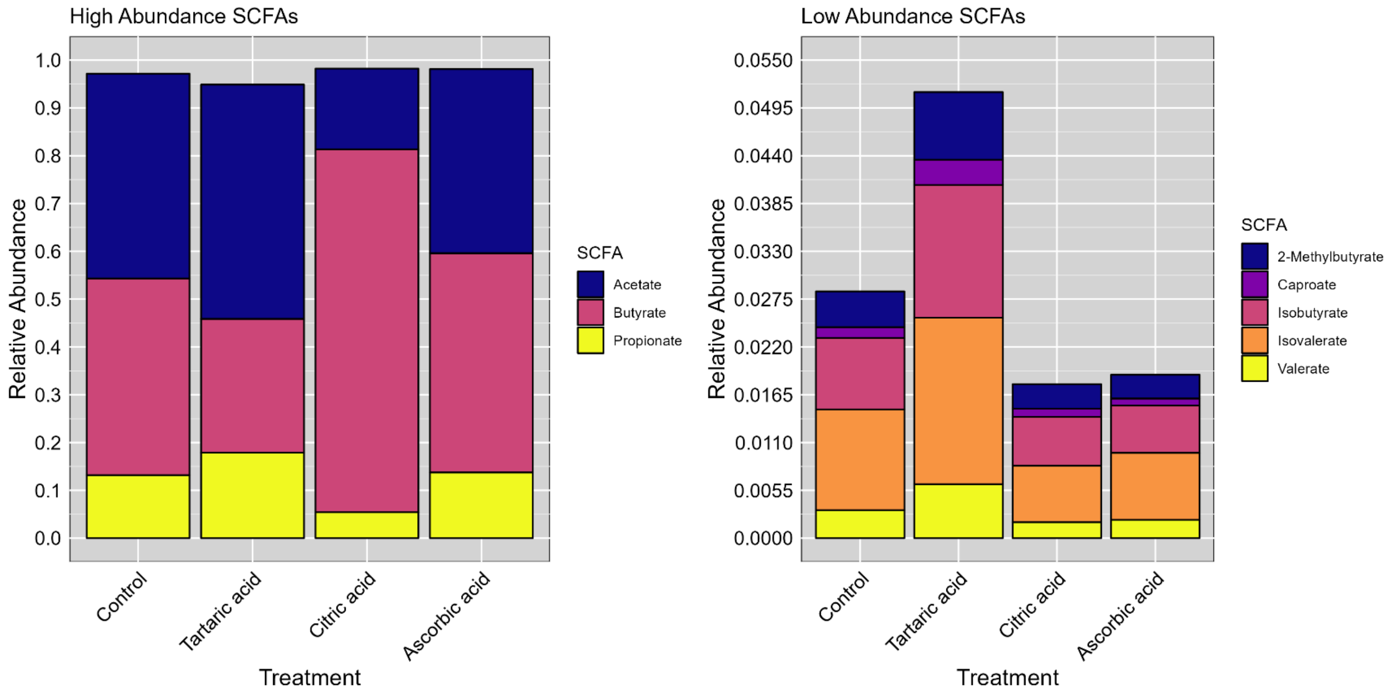

C

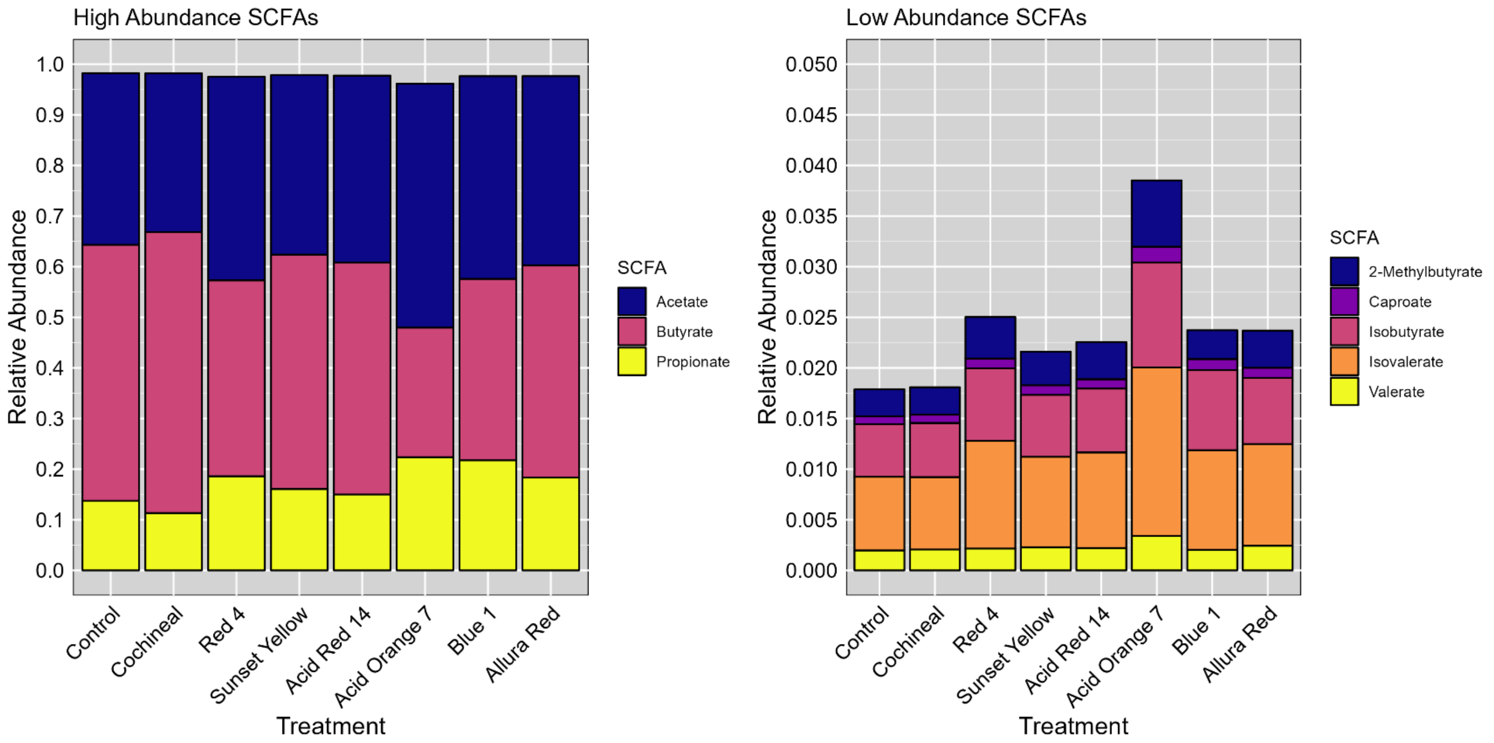
High Abundance SCFAs
Relative Abundance
1.0
0.9
0.8
0.7
0.6
0.5
0.4
0.3
0.2
0.1
0.0
Control
Cochineal
Red 4
Sunset Yellow
Acid Red 14
Acid Orange 7
Blue 1
Allura Red
Treatment
SCFA
Acetate
Butyrate
Propionate
Low Abundance SCFAs
Relative Abundance
0.050
0.045
0.040
0.035
0.030
0.025
0.020
0.015
0.010
0.005
0.000
Control
Cochineal
Red 4
Sunset Yellow
Acid Red 14
Acid Orange 7
Blue 1
Allura Red
Treatment
SCFA
2-Methylbutyrate
Caproate
Isobutyrate
Isovalerate
Valerate

D

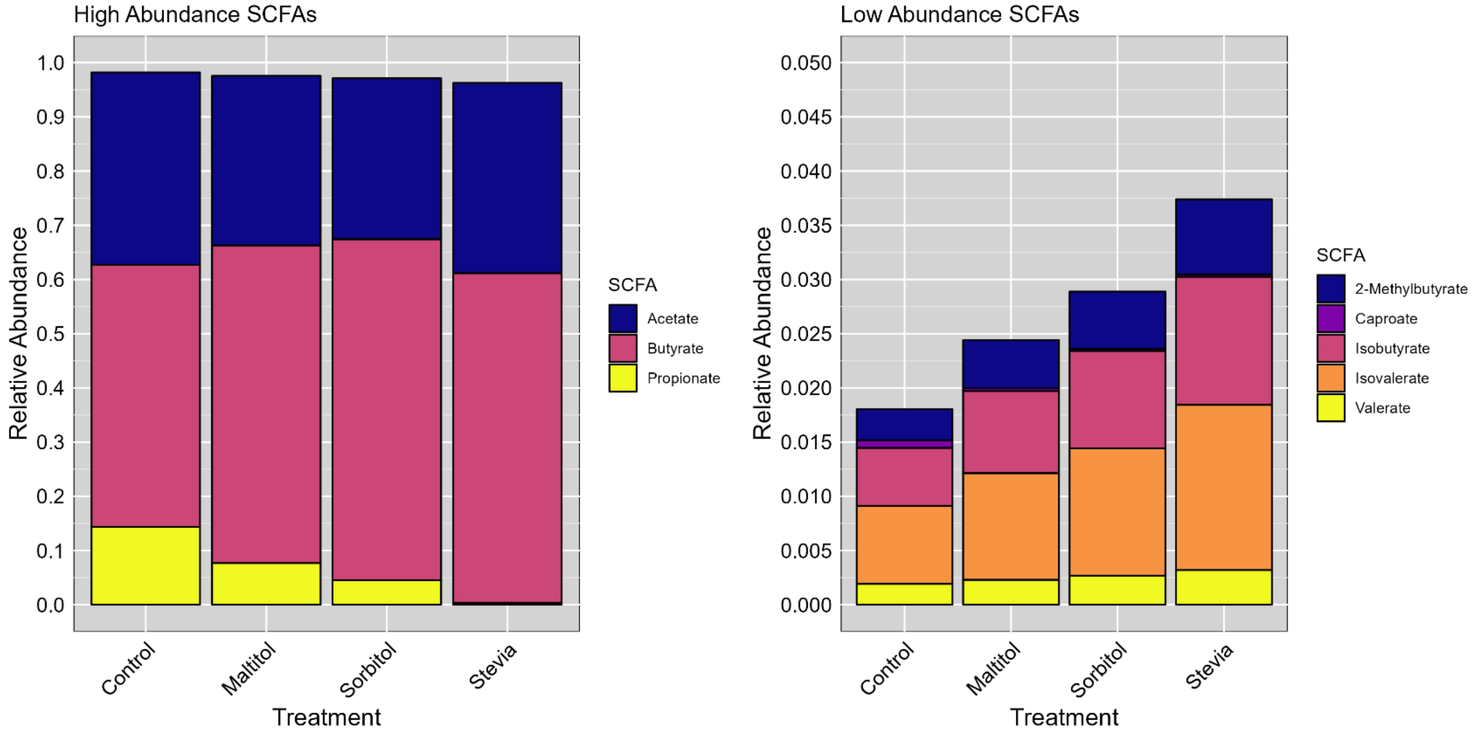
High Abundance SCFAs
Relative Abundance
1.0
0.9
0.8
0.7
0.6
0.5
0.4
0.3
0.2
0.1
0.0
Control
Maltitol
Sorbitol
Stevia
Treatment
SCFA
Acetate
Butyrate
Propionate
Low Abundance SCFAs
Relative Abundance
0.050
0.045
0.040
0.035
0.030
0.025
0.020
0.015
0.010
0.005
0.000
Control
Maltitol
Sorbitol
Stevia
Treatment
SCFA
2-Methylbutyrate
Caproate
Isobutyrate
Isovalerate
Valerate
