## Supplementary material for "High-Throughput Screening of the Effects of 90 Xenobiotics on the Simplified Human Gut Microbiota Model (SIHUMIx): A Metaproteomic and Metabolomic Study": Figure S5

**Figure S5: Impact of food additives, and dyes on the metabolic pathways of SIHUMIx.** The pathway intensity was measured by metaproteomics and is displayed as $log_2FC$ for food additives and dyes which affected at least the intensity of one pathway compared to the control. Statistically significant effects ($P_{adj}$ <0.05) are highlighted by asterisks

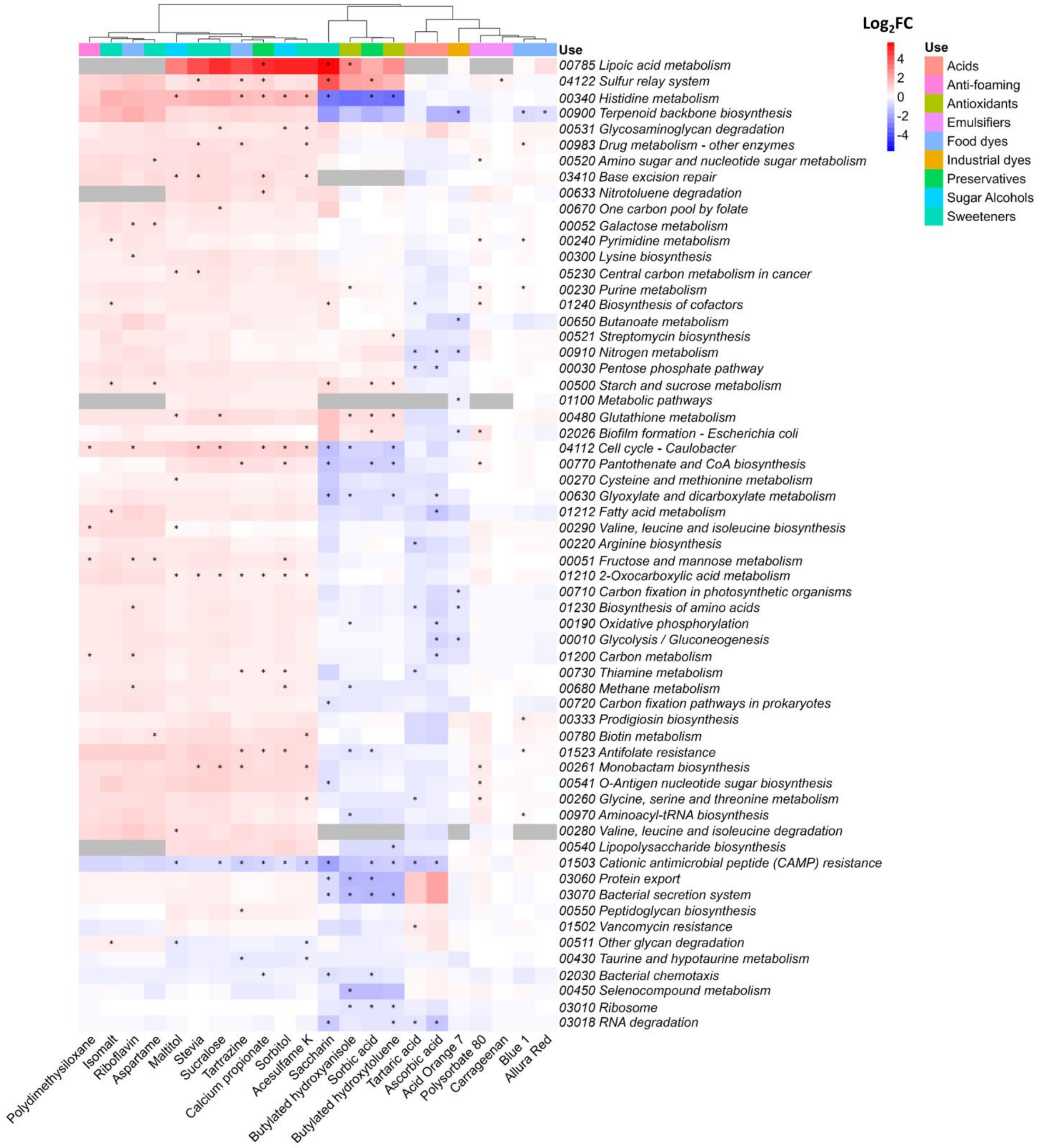
