## Supplementary material for "High-Throughput Screening of the Effects of 90 Xenobiotics on the Simplified Human Gut Microbiota Model (SIHUMIx): A Metaproteomic and Metabolomic Study": Table S1

**Table S1: Complex intestinal medium (CIM) composition**

| **Ingredient** | **Quantity [g/L]** | **supplier** |
| --- | --- | --- |
| Arabinogalactan (larch wood) | 2 | Sigma-Aldrich |
| Bile Acids sodium salt | 0.5 | Sigma-Aldrich |
| Calcium chloride x 2 H_2_O | 0.01 | Merck |
| Casein peptone (pancreatic) | 4.3 | Roth |
| Di-Potassium hydrogen phosphate | 0.04 | Roth |
| Hemin (bovine) | 0.005 | Sigma-Aldrich |
| Inulin | 1 | Serva |
| L-cysteine hydrochloride | 0.5 | Biochemica |
| Magnesium sulfate | 0.01 | Roth |
| Menadione | 0.001 | Sigma-Aldrich |
| Mucin (porcine gastric Type II) | 4 | Sigma-Aldrich |
| Pectin, citrus peel | 2 | Sigma-Aldrich |
| Potassium di-hydrogen phosphate | 0.04 | Roth |
| Sodium chloride | 0.72 | Roth |
| Sodium hydrogen carbonate | 2 | Roth |
| Starch, wheat | 5 | Roth |
| Xylo-oligosaccharide (corn) | 2 | Roth |
| Yeast extract | 2 | Chemsolut |
